# Show me your secret(ed) weapons: a multifaceted approach reveals novel type III-secreted effectors of a plant pathogenic bacterium

**DOI:** 10.1101/679126

**Authors:** Irene Jiménez Guerrero, Francisco Pérez-Montaño, Gustavo Mateus da Silva, Naama Wagner, Dafna Shkedy, Mei Zhao, Lorena Pizarro, Maya Bar, Ron Walcott, Guido Sessa, Tal Pupko, Saul Burdman

## Abstract

Many Gram-negative plant and animal pathogenic bacteria employ a type III secretion system (T3SS) to secrete protein effectors into the cells of their hosts and promote disease. The plant pathogen *Acidovorax citrulli* requires a functional T3SS for pathogenicity. As with *Xanthomonas* and *Ralstonia spp*., an AraC-type transcriptional regulator, HrpX, regulates expression of genes encoding T3SS components and type III-secreted effectors (T3Es) in *A. citrulli*. A previous study reported eleven T3E genes in this pathogen, based on the annotation of a sequenced strain. We hypothesized that this was an underestimation. Guided by this hypothesis, we aimed at uncovering the T3E arsenal of the *A. citrulli* model strain, M6. We carried out a thorough sequence analysis searching for similarity to known T3Es from other bacteria. This analysis revealed 51 *A. citrulli* genes whose products are similar to known T3Es. Further, we combined machine learning and transcriptomics to identify novel T3Es. The machine learning approach ranked all *A. citrulli* M6 genes according to their propensity to encode T3Es. RNA-Seq revealed differential gene expression between wild-type M6 and a mutant defective in HrpX. Data combined from these approaches led to the identification of seven novel T3E candidates, that were further validated using a T3SS-dependent translocation assay. These T3E genes encode hypothetical proteins, do not show any similarity to known effectors from other bacteria, and seem to be restricted to plant pathogenic *Acidovorax* species. Transient expression in *Nicotiana benthamiana* revealed that two of these T3Es localize to the cell nucleus and one interacts with the endoplasmic reticulum. This study not only uncovered the arsenal of T3Es of an important pathogen, but it also places *A. citrulli* among the “richest” bacterial pathogens in terms of T3E cargo. It also revealed novel T3Es that appear to be involved in the pathoadaptive evolution of plant pathogenic *Acidovorax* species.

**Author summary:** *Acidovorax citrulli* is a Gram-negative bacterium that causes bacterial fruit blotch (BFB) disease of cucurbits. This disease represents a serious threat to cucurbit crop production worldwide. Despite the agricultural importance of BFB, the knowledge about basic aspects of *A*. citrulli-plant interactions is rather limited. As many Gram-negative plant and animal pathogenic bacteria, *A. citrulli* employs a complex secretion system, named type III secretion system, to deliver protein virulence effectors into the host cells. In this work we aimed at uncovering the arsenal of type III-secreted effectors (T3Es) of this pathogen by combination of bioinformatics and experimental approaches. We found that this bacterium possesses at least 51 genes that are similar to T3E genes from other pathogenic bacteria. In addition, our study revealed seven novel T3Es that seem to occur only in *A. citrulli* strains and in other plant pathogenic *Acidovorax* species. We found that two of these T3Es localize to the plant cell nucleus while one partially interacts with the endoplasmic reticulum. Further characterization of the novel T3Es identified in this study may uncover new host targets of pathogen effectors and new mechanisms by which pathogenic bacteria manipulate their hosts.

## Introduction

The genus *Acidovorax* (class Betaproteobacteria) contains a variety of species with different lifestyles. While some species are well adapted to soil and water environments, others have developed intimate relationships with eukaryotic organisms, including as plant pathogens [1]. Among the latter, *Acidovorax citrulli* is one of the most important plant pathogenic species [2]. This bacterium infects all aerial parts of cucurbit plants, causing bacterial fruit blotch (BFB) disease. The unavailability of effective tools for managing BFB, including the lack of resistance sources, and the disease’s high destructive potential, exacerbate the threat BFB poses to cucurbit (mainly melon and watermelon) production [3, 4]. Despite the economic importance of BFB, little is known about basic aspects of *A*. citrulli-plant interactions.

On the basis of genetic and biochemical features, *A. citrulli* strains are divided into two main groups: group I strains have been generally isolated from melon and other non-watermelon cucurbits, whereas group II strains have been mainly isolated from watermelon [5–7]. *Acidovorax citrulli* M6 is a group I strain that was isolated in 2002 from a BFB outbreak of melons in Israel [5], and subsequently became a model group I strain for investigation of basic aspects of BFB. The *A. citrulli* M6 genome has been sequenced, first by Illumina MiSeq [8] and recently, by PacBio [9], which allowed its complete closure.

As many Gram-negative plant and animal pathogenic bacteria, *A. citrulli* relies on a functional type III secretion system (T3SS) to promote disease [10]. This complex secretion system is employed by these pathogens to deliver protein effectors into target eukaryotic cells. Collectively, type III-secreted effectors (T3Es) promote disease by modulating a variety of cellular functions for the benefit of the pathogen [11–13]. In the case of plant pathogenic bacteria, type III-secreted effectors (T3Es) were shown to promote virulence through alteration of the plant cell metabolism and/or suppression of host immune responses [14, 15]. As part of their defence mechanism, plants recognize some effectors by corresponding disease resistance (R) proteins, mostly belonging to the nucleotide-binding (NB)-leucine-rich repeat (LRR) type of immune receptors (NLRs) [16, 17]. Upon effector recognition, the R protein elicits a battery of defense responses collectively referred to as effector-triggered immunity (ETI). ETI is often accompanied by the hypersensitive response (HR), a rapid death of plant cells at the infection site that arrests pathogen spread in the plant tissue [18]. Therefore, elucidating the arsenal of effectors and their contribution to virulence, are of critical importance for the understanding of basic aspects of pathogenicity but also for translational research in the crop protection field.

Due to the requirement of type III secretion (T3S) for pathogenicity in susceptible plants and HR elicitation in resistant plants, the genes encoding key T3SS regulators and structural components in plant pathogenic bacteria are named *hrp* genes (for <u>HR</u> and <u>p</u>athogenicity) or *<u>hrc</u>* genes, in the <u>c</u>ase of *hrp* genes that are conserved among different bacterial genera, including in animal pathogens [19]. On the basis of gene content, operon organization and regulation, *hrp* clusters are divided into two classes: class I contains the *hrp* clusters of *Pseudomonas syringae* and enteric plant pathogenic bacteria, while class II contains the clusters of *Xanthomonas* species, *Ralstonia solanacearum* and plant pathogenic *Acidovorax* spp. [10, 19, 20].

In *Xanthomonas* spp. and *R. solanacearum*, the expression of *hrp, hrc* and *<u>h</u>r<u>p</u>*-<u>a</u>ssociated (hpa) genes, as well as of some T3E genes, is regulated by HrpG and HrpX/HrpB (HrpX in *Xanthomonas* spp. and HrpB in *R. solanacearum)*. HrpG belongs to the OmpR family of two-component system response regulators and controls expression of *hrpX/hrpB* [21–23]. *hrpX* and *hrpB* encode AraC-type transcriptional activators that directly mediate the expression of most *hrp/hrc* operons and many T3E genes, via binding to DNA motifs that are present in their promoter regions. These DNA motifs are named plant-inducible promoter (PIP) box (TTCGB-N15-TTCGB; B being any nucleotide except adenine) in *Xanthomonas* spp. [24] and hrp_II_ box (TTCG-N16_TTCG) in *R. solanacearum* [25]. Recently, Zhang *et al*. showed that the *hrpG* and *hrpX/hrpB* (thereafter *hrpX)* orthologous genes of the *A. citrulli* group II strain Aac5 are required for pathogenicity [26]. They also showed that HrpG activates expression of *hrpX*, which in turn, regulates the expression of a T3E gene belonging to the YopJ family.

Until recently, based on the annotation of the genome of the *A. citrulli* group II strain AAC00-1, we were aware of eleven genes showing similarity to known T3E genes from other bacteria [27]. Considering the higher numbers of T3E genes in several other plant pathogenic bacteria, we hypothesized that this is an underestimation of the actual number of T3Es in *A. citrulli*. We also hypothesized that *A. citrulli* may carry novel T3E genes that were not previously described in other bacteria. Guided by these hypotheses, we carried out a detailed sequence analysis of *A. citrulli* M6 open reading frames (ORFs) to identify genes with similarity to known T3E genes from other bacteria. We also combined machine-learning (ML) and RNA-Seq approaches to identify putative, novel *A. citrulli* T3Es. Further, we adapted a T3E translocation assay to verify T3S-dependent translocation of candidate effectors. Combining these approaches allowed identification of seven new T3Es that appear to be unique to plant pathogenic *Acidovorax* species. Subcellular localization of three of these T3Es in *N. benthamiana* leaves was also determined by *Agrobacterium*-mediated transient expression.

## Results

### Identification of new T3E genes of *A. citrulli* by genome annotation, machine learning and sequence analyses

Analysis of the genome of the group II *A. citrulli* strain AAC00-1 (GenBank accession CP000512.1) revealed eleven genes similar to T3E genes of other plant pathogenic bacteria [27]. These genes were present in all tested group II strains. In contrast, all assessed group I strains, including M6, lacked the effector gene *Aave_2708* (gene ID according to the AAC00-1 annotation), encoding a *Xanthomonas euvesicatoria* XopJ homolog. Group I strains also had disrupted open reading frames (ORFs) in the genes *Aave_3062*, encoding an effector similar to *Xanthomonas oryzae* pv. *oryzicola* AvrRxo1, and *Aave_2166*, encoding a *X. euvesicatoria* AvrBsT homolog [27].

To identify new putative T3E genes of *A. citrulli* we applied a machine learning (ML) approach, that was successfully utilized for identification of new T3E genes of *X. euvesicatoria* [28] and *Pantoea agglomerans* [29]. Using this algorithm, all ORFs of a bacterial genome are scored according to their propensity to encode T3Es. The scoring is based on a large set of features including similarity to known T3E genes, genomic organization, amino acid composition bias, characteristics of the putative N-terminal translocation signal and GC content, among others (see Methods).

An initial ML run was used to classify all ORFs of strain AAC00-1 according to their probability to encode T3Es. This strain, rather than M6, was used for learning and prediction, because at the time this ML was conducted, the AAC00-1 genome was fully assembled with better annotation. For training, the positive set included 12 AAC00-1 genes that encoded T3E homologs: the eleven genes described by Eckshtain-Levi *et al*. [27] and one additional gene, *Aave_2938* that is identical to *Aave_2708*. The negative set included genes that showed high sequence similarity to ORFs of a non-pathogenic *Escherichia coli* strain. The output of this ML run was a list of all annotated genes of *A. citrulli* AAC00-1 ranked by their propensity to encode T3Es (S1 Table). For each ORF, we searched for the homolog in *A. citrulli* M6. Among the top predictions from AAC00-1, many genes did not have homologs in M6. As expected, the aforementioned 12 positive T3E genes of AAC00-1 were ranked high in this list (among the 36 highest scoring predictions, with eight being ranked among the top 10, and eleven among the top 15; S1 Table). Results from this first ML run served, together with RNA-Seq data, as the basis for selection of candidate T3E (CT3E) genes for experimental validation (see below).

In parallel, we performed an extensive homology search, using BlastP, to identify additional putative T3E genes of *A. citrulli* M6. This analysis led to the identification of many additional genes with significant similarity to T3E genes from other plant pathogenic bacteria. Table 1 summarizes the arsenal of putative T3E genes of *A. citrulli* M6, based on its genome annotation and sequence similarity analysis. Overall, we found 51 putative T3E genes in the *A. citrulli* M6 genome, in support of the notion that *A. citrulli* has a larger T3E repertoire than previously estimated. Most of these genes also received high scores in the ML search ranking among the top 100 ORFs (Table 1 and S1 Table). With that said, ten genes encoding T3E homologs were ranked in very low positions in the ML run (positions 231 to 1161; Table 1). On the other hand, many top ranked genes were annotated as encoding hypothetical proteins, some of which could encode yet unknown T3Es.

**Table 1.**
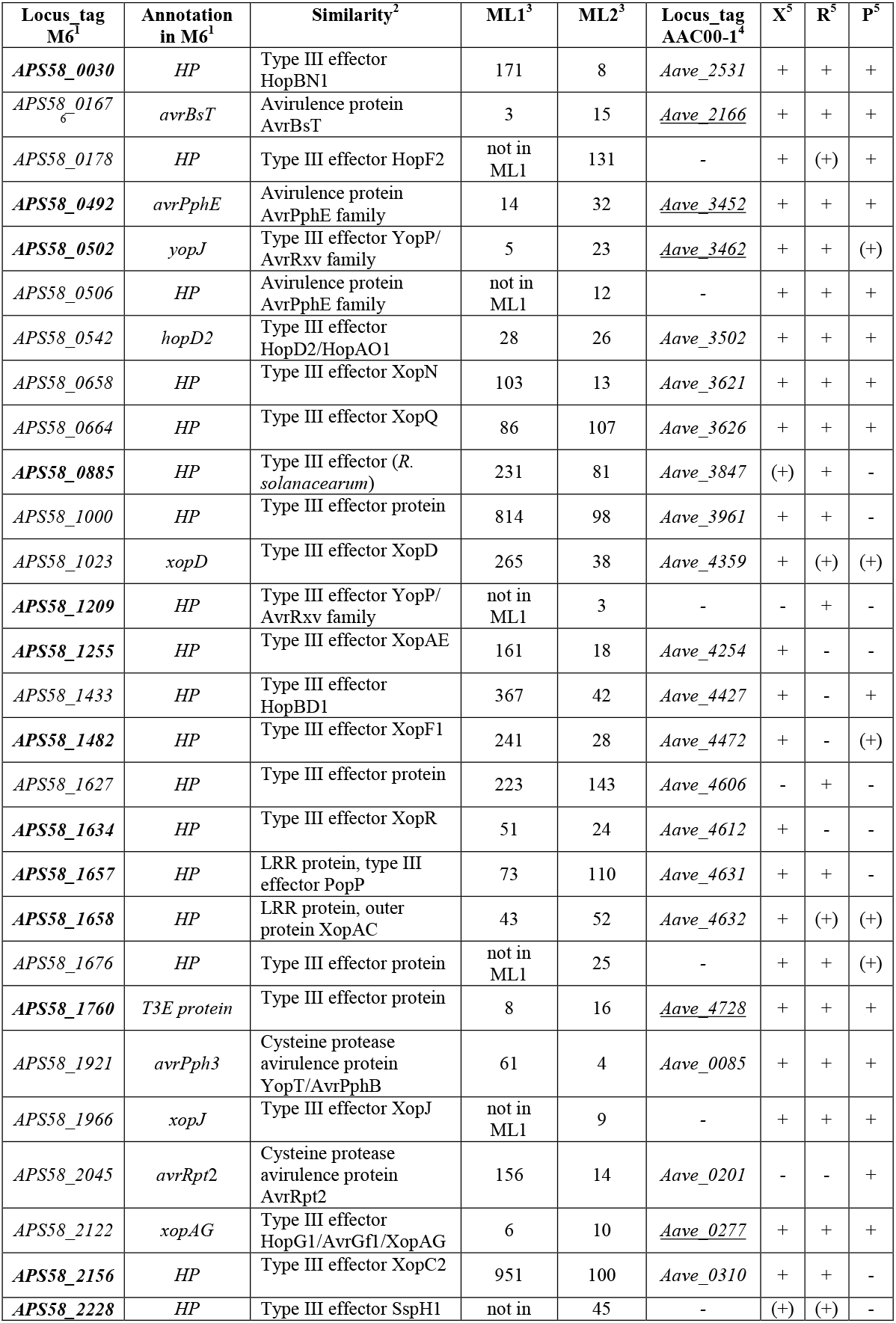
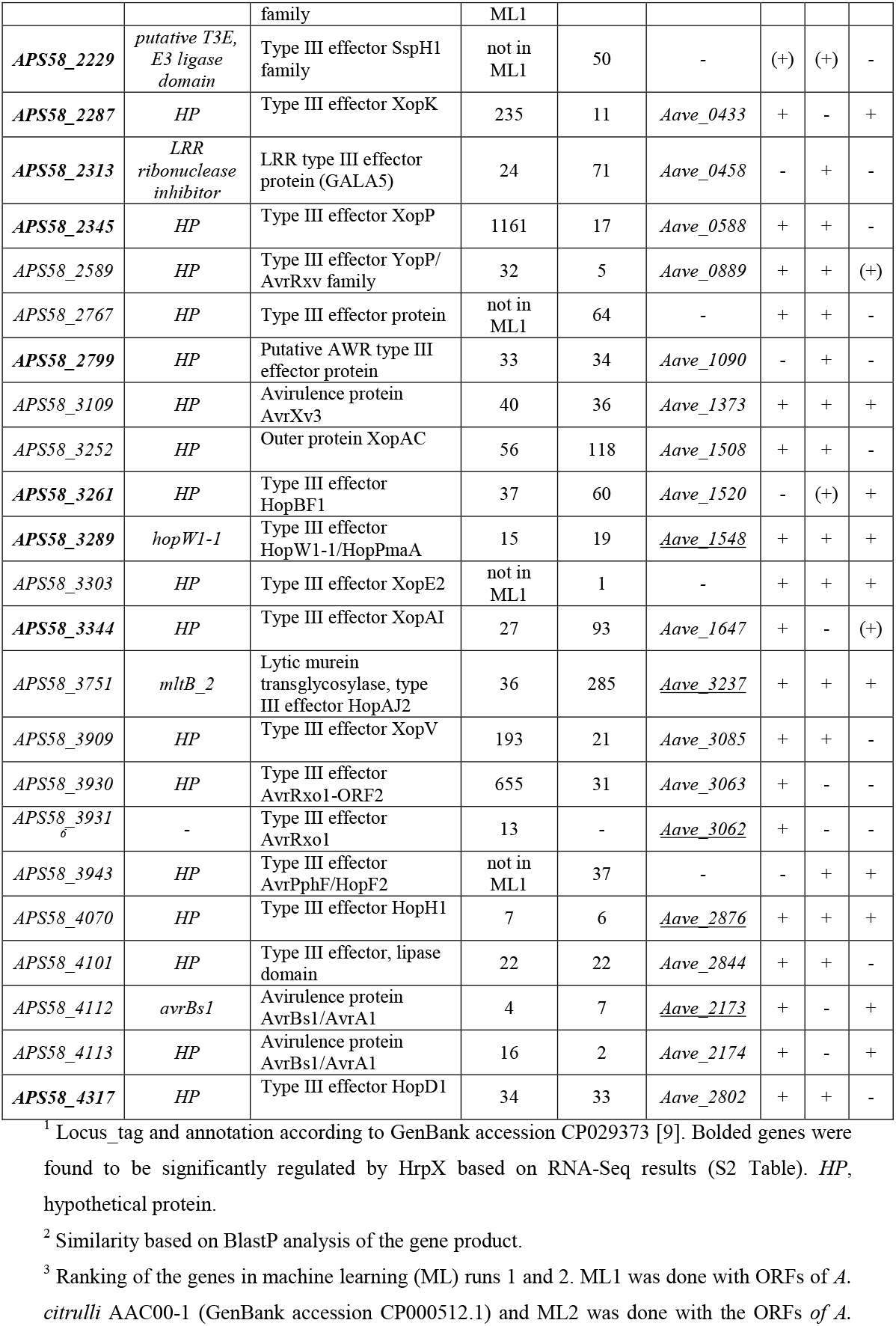
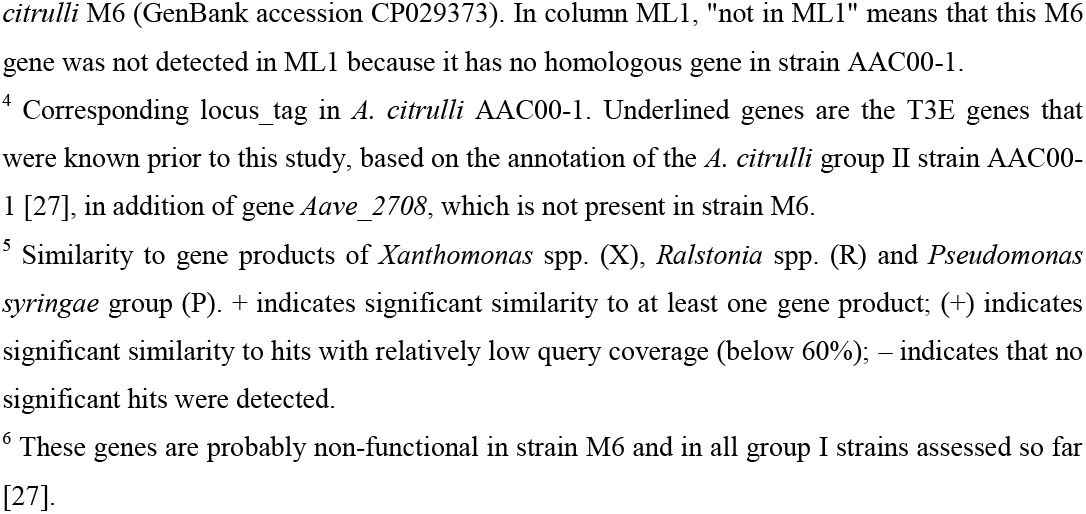
List of putative T3E genes of *Acidovorax citrulli* M6 based on genome annotation and sequence similarity (BlastP) to known T3E genes from other plant pathogenic bacteria.

An additional insight of this analysis was that most predicted T3E genes of *A. citrulli* share levels of similarity with T3E genes of *Xanthomonas* spp. and *R. solanacearum* (41 and 40 genes, respectively; Table 1). A smaller number of genes, 31, shared similarity with T3E genes of *P. syringae* strains. We also assessed the occurrence of these T3Es in other plant pathogenic *Acidovorax* species (S2 Table). Except for the HopBD1 homolog APS58_1433 that could be detected only in *A. citrulli* strains, the other predicted T3Es occur in other pathogenic *Acidovorax* species, with some of them being widely distributed. For instance, the putative effectors APS58_0492, APS58_0506, APS58_1482, APS58_1657, APS58_1658, APS58_2228, APS58_2313, APS58_2345, APS58_2799, APS58_3303 and APS58_3751 could be detected, at different levels of similarity, in all species. The other putative effectors were restricted to fewer species, with most of them being detected in *A. avenae* strains. While this may reflect the close relatedness between *A. citrulli* and *A. avenae* [30], it is important to consider that, at the time of this analysis, the public database contained 7 and 18 genomes of *A. citrulli* and *A. avenae* strains, respectively, but only two draft genomes of *A. oryzae* and one draft genome for each of the other species.

Interestingly, of the 51 putative T3E genes of *A. citrulli* M6, ten were not present in the genome of the group II strain AAC00-1 (Table 1). Besides M6 and AAC00-1, the NCBI database includes draft genomes of one additional group II strain, KAAC17055, and four group I strains (pslb65, tw6, DSM 17060 and ZJU1106). BlastN analyses revealed that these ten genes are also absent in strain KAAC17055, but present in most of the group I strains. The only exceptions were *APS58 0506* that was not detected in strains tw6 and DSM 17060, *APS58 1209* that was not detected in tw6, and *APS58 2767* that was not detected in DSM 17060. The inability to detect these T3E genes in the genomes of strains tw6 and DSM 17060 could reflect true absence in these strains but also could be due to the draft nature of these genomes. In any case, these results strongly suggest that the ten M6 T3E genes that are absent in the group II strains AAC00-1 and KAAC17055 could be specific to group I strains of *A. citrulli*. Yet, this assumption should be verified on a larger collection of strains. Interestingly, among these ten T3E genes, *APS58 0506, APS58 2228* and *APS58 3303*, were detected in strains of all other plant pathogenic *Acidovorax* species (S2 Table). In the case of *APS58 2228*, it should be mentioned that the group II strains AAC00-1 and KAAC17055 possess genes *(Aave_0378* in AAC00-1) that encode short products (140 a. a.) and partially align with the C-terminal region of the group I product (with predicted length of 538 a.a.). In our analysis we did not consider them as ortholog genes.

### HrpX is required for pathogenicity of *A. citrulli* M6 and regulates the expression of T3SS components and T3E genes

In *Xanthomonas* spp. and *R. solanacearum*, the transcriptional regulator HrpX (HrpB in *R. solanacearum)* plays a key role in regulation of *hrp* and T3E genes. We hypothesized that this is also the case in *A. citrulli* M6. To assess this hypothesis, we first generated an *A. citrulli* M6 strain mutated in *APS582298*, the *hrpX* orthologous gene. This mutant lost the ability to cause disease in melon (Fig 1A) and induce HR in pepper leaves (Fig 1B), as previously observed for a strain carrying a mutation in the *hrcV* gene, which encodes a core component of the T3SS [10]. A similar loss of pathogenicity was observed for a mutant defected in the *hrpG* homolog gene, *APS58_2299* (S1 Fig). Complementation of both *hrpX* and *hrpG* mutations restored pathogenicity, although necrotic symptoms induced by the complemented strains were less severe than those induced by the wild-type strain (S1 Fig).

**Fig 1.**
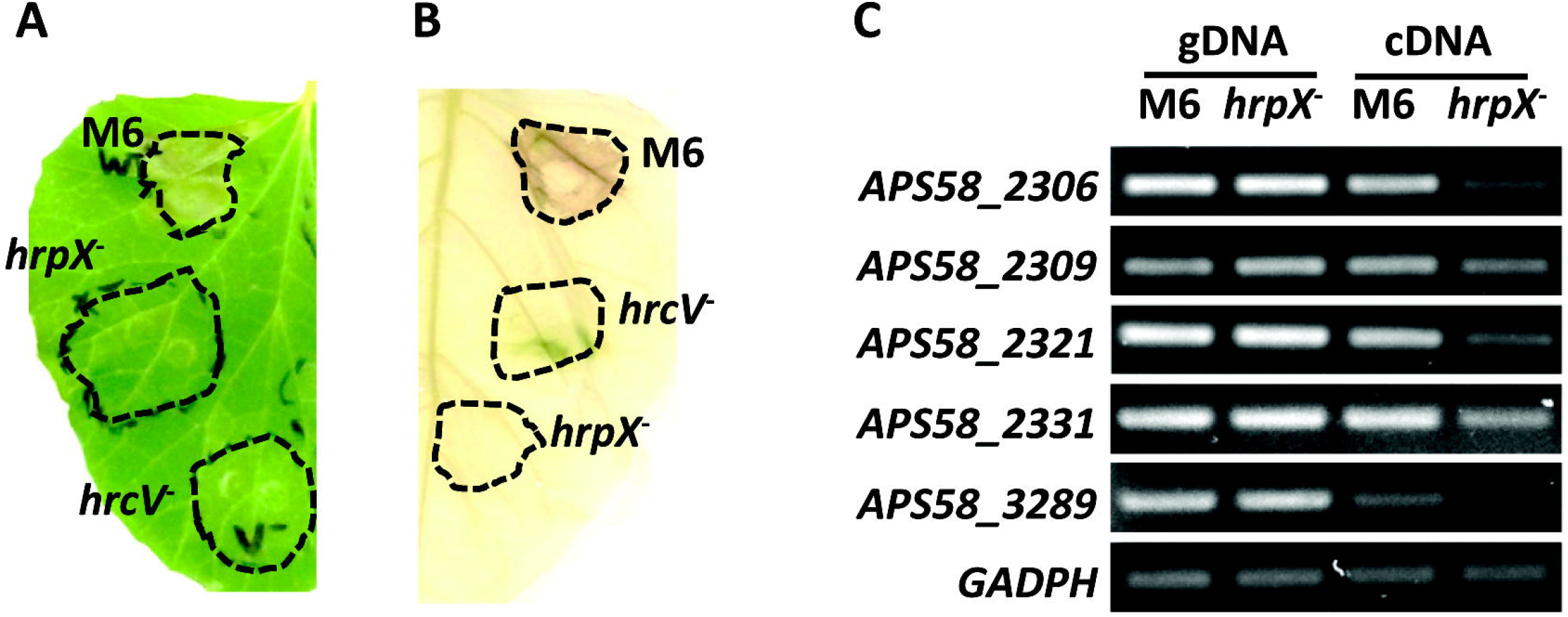
HrpX is required for pathogenicity and regulates expression of T3S and T3E genes in *Acidovorax citrulli* M6. (A) Disease lesions produced in a melon leaf inoculated with wild-type M6, but not with mutant strains defective in *hrpX* or *hrcV* (encoding a core component of the T3SS) genes. The picture was taken at 3 days after infiltration (d.a.i.). (B) Cell death observed in a pepper leaf following inoculation with wild-type M6, but not with *hrpX* and *hrcV* mutants. The picture was taken at 4 d.a.i. In (A) and (B), leaves were syringe-infiltrated with bacterial suspension of 10^8^ CFU/ml. (C) Qualitative assessment of differential gene expression between wild-type M6 and the M6 *hrpX* mutant after 72 h of growth in XVM2 minimal medium at 28 °C. gDNA, amplification of genomic DNA. cDNA, reverse-trancriptase (RT)-PCR of RNA extracts. Genes: *hrcV (APS58_2306), hrcT (APS58_2309), hrcJ (APS58_2321)* and *hrcC (APS58_2331)*, encoding core components of the T3SS; *APS58_3289*, encoding a T3E similar to *Pseudomonas syringae hopW1-1*; and *GADPH*, glyceraldehyde-3-phosphate dehydrogenase *(APS58_1610;* control gene).

Further, we used reverse transcription-PCR (RT-PCR) to compare expression of four genes encoding T3SS components and one T3E gene *(APS58_3289*, encoding a *P. syringae hopW1-1* homolog of *hopW1-1)* between the *hrpX* mutant and wild-type M6 following growth in XVM2 medium. This medium was optimized for expression of T3S genes in *X. euvesicatoria*, as it simulates, to some extent, the plant apoplast environment [31]. After 72 h of growth, expression of the tested genes was reduced in the *hrpX* mutant relative to wild-type M6 (Fig 1C).

### Identification of HrpX-regulated genes by RNA-Seq

Based on RT-PCR results, we carried out RNA-Seq analysis to compare gene expression between wild-type M6 and the *hrpX* mutant, after 72 h of growth in XVM2 medium. This approach revealed 187 genes showing significant differential expression (significant fold-change of ± 2) between the strains (Fig 2A). Of these, 159 genes had significantly reduced expression in the *hrpX* mutant relative to wild-type M6, while 28 genes showed the opposite pattern (S3A and S3B Tables). RNA-Seq results were validated by qPCR experiments that confirmed lower expression of 10 tested genes in the *hrpX* mutant under the same conditions (Fig 2B).

**Fig. 2.**
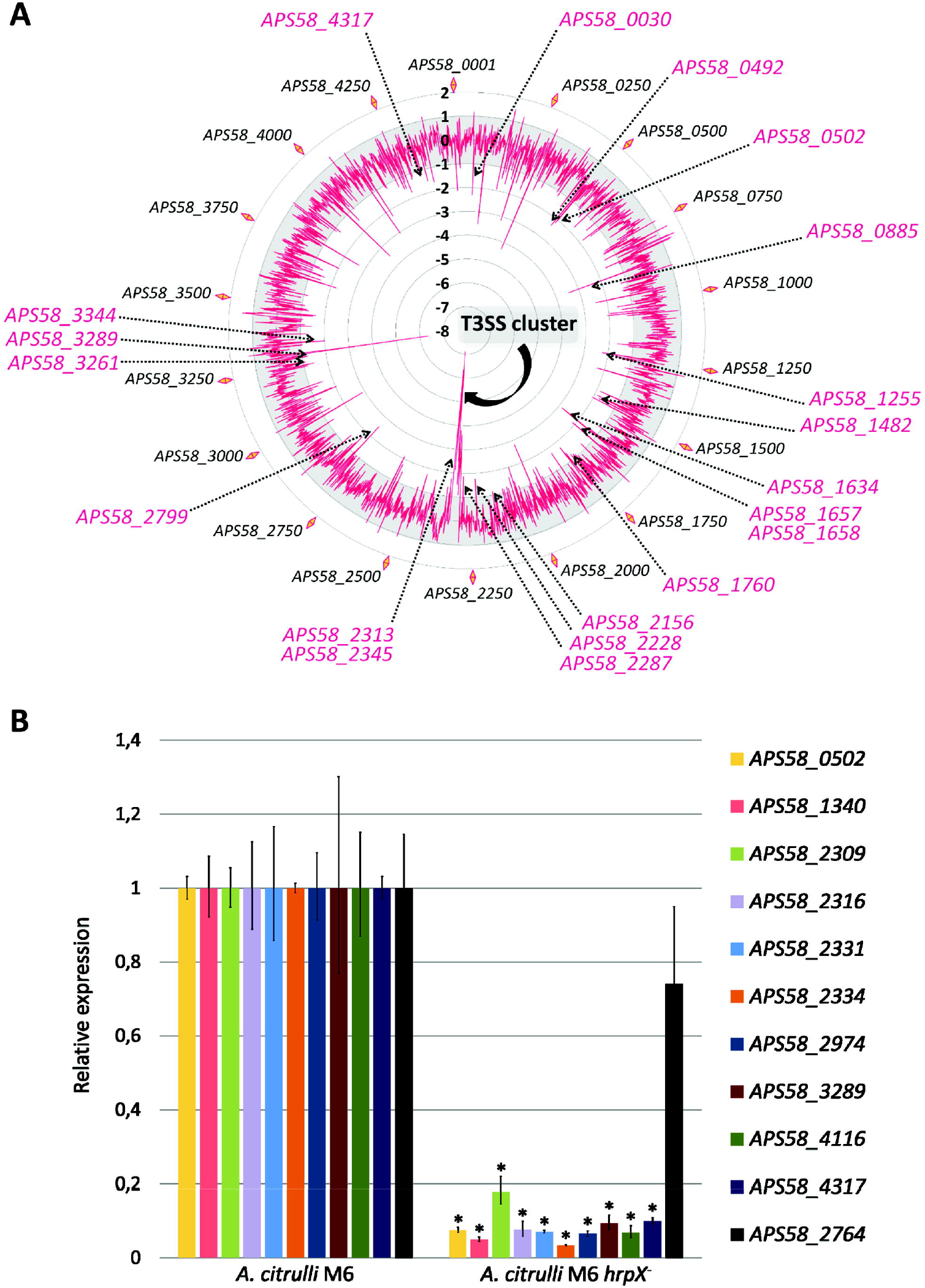
Comparative transcriptomics analysis between *Acidovorax citrulli* M6 and the M6 *hrpX* mutant. (A) Relative gene expression profile as assessed by RNA-Seq of cells grown for 72 h at 28 °C in minimal XVM2 medium (2 and 3 replicates for the *hrpX* mutant and wild-type strain, respectively). The *A. citrulli* M6 genome map is represented in the external circle. Internal red line shows differential gene expression between the strains. Genes within the gray zone: no significant differences between the strains. The −8 to 2 scale indicates relative expression of the mutant compared with the wild-type. Genes with significantly reduced or increased expression in the mutant relative to the wild-type strain are in the inner and outer regions relative to the gray zone, respectively. Arrows indicate the Hrp-T3SS cluster as well as genes with homology to known effectors from other plant pathogenic bacteria. (B) Relative expression of selected genes by qRT-PCR following bacterial growth under identical conditions as for the RNA-Seq experiment (3 biological replicates per strain). Asterisks indicate significant differences between wild-type and *hrpX* mutant at α = 5% by the Mann-Whitney non-parametrical test. All tested genes except *APS58_2764* showed significantly reduced expression in the mutant relative to strain M6 in the RNA-Seq analysis.

Most HrpX-regulated genes could not be assigned to Gene Ontology (GO) categories using Blast2GO. Of the 159 genes that showed reduced expression in the *hrpX* mutant, only 47 were assigned to at least one biological process category. Blast2GO results are detailed in S3C and S3D Tables, and Fig 3 shows the number of biological process categories of genes with reduced expression in the mutant. Among the most frequent categories, 10 hits were found for transmembrane transport proteins, including several ABC transporters and permeases, and 6 matched with regulation of transcription. Nine hits belonged to protein secretion/protein secretion by the T3SS and these corresponded to genes encoding Hrp/Hrc components. Notably, most T3S and T3E genes could not be assigned to any specific GO biological process; this was the case for 11 *hrp/hrc/hpa* genes and for 24 T3E genes (S3C Table). Overall, RNA-Seq revealed 20 *hrp*/*hrc/hpa* genes and 27 genes encoding putative T3Es (including the seven new effectors identified in this study; see below) that had significantly reduced expression in the *hrpX* mutant relative to wild-type M6 (S3B and S3C Tables). Importantly, almost 60 genes that showed reduced expression in the *hrpX* mutant are annotated as hypothetical proteins and did not show similarity to known T3E genes. It is possible that some of these genes encode novel T3Es.

**Fig. 3.**
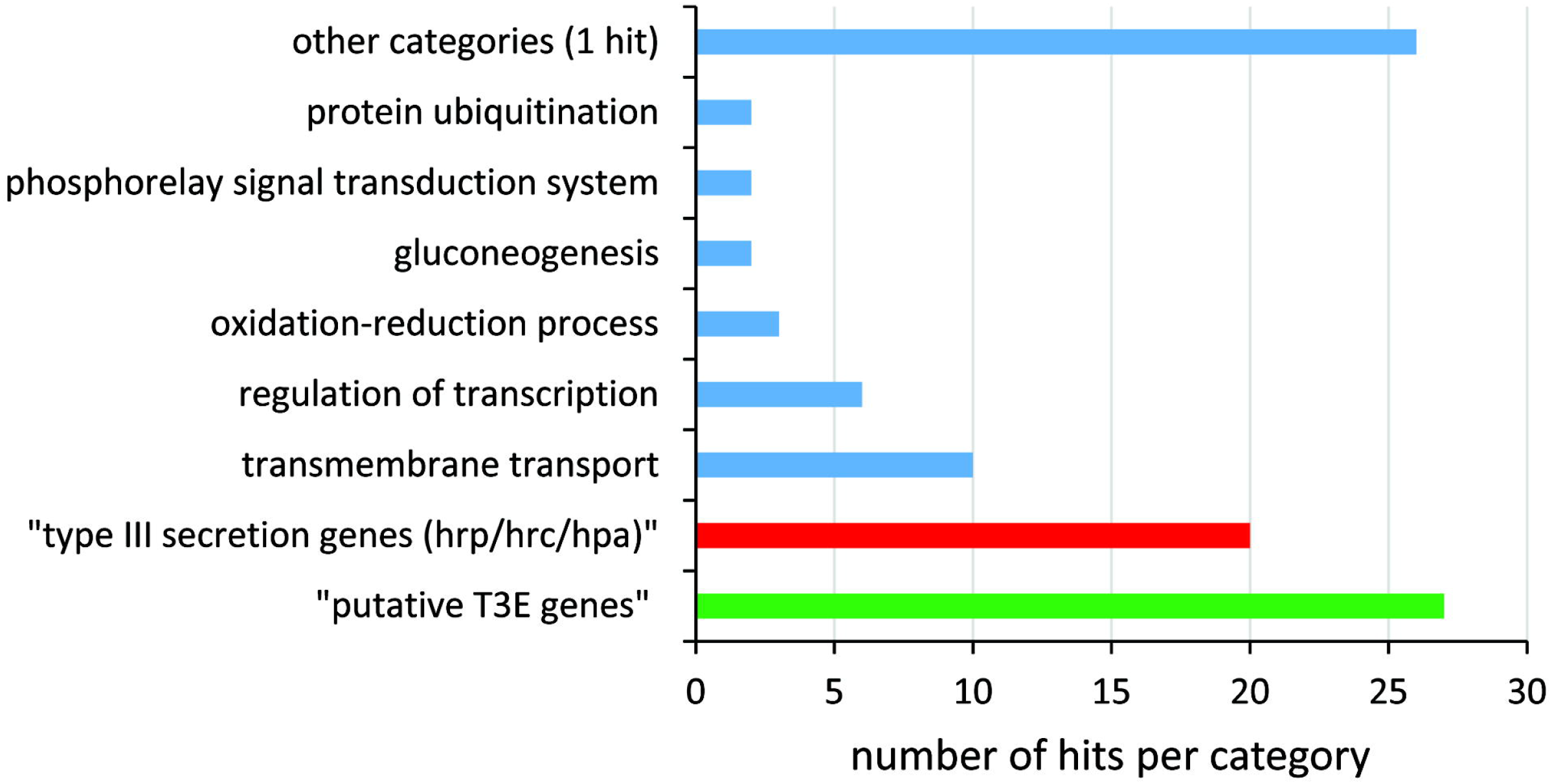
Distribution of *Acidovorax citrulli* M6 HrpX-regulated genes among categories of biological processes. Of the 159 genes that showed reduced expression in the *hrpX* mutant relative to wild-type M6, only 47 could be assigned to at least one Gene ontology (GO) biological process category (blue columns). HrpX-regulated genes encoding T3S structural and accessory proteins (red column) and putative T3Es (green column) were manually assigned to these categories.

Interestingly, the *hrpX* mutant also showed reduced expression of several genes encoding proteins that are putatively secreted by the type II secretion system (T2SS). We used SignalP, Pred-Tat and Phobius tools to detect putative Tat or Sec type II secretion (T2S) signals in the ORFs of all genes that showed significantly lower expression in the *hrpX* mutant relative to the wild-type strain. While T2S signals were predicted in 39 genes by at least one of the tools (not shown), 14 genes were predicted to encode products with T2S signals by the three different tools (S3E Table). Among these genes were *APS58 0633 (xynB)* encoding 1-4-β-xylanase, *APS58_2599 (pelA_2)*, encoding pectate lyase, and *APS58_3 722*, encoding a family S1 extracellular serine protease. These three genes were also shown to contain PIP boxes in their promoter region (S3B Table).

Of the 28 genes showing increased expression in the *hrpX* mutant relative to wild-type M6, only ten could be assigned to GO categories, most of which belonged to regulatory genes (regulation of transcription, phosphorelay signal transduction system, signal transduction; S3D Table).

### Identification of PIP boxes in HrpX-regulated genes

We used fuzznuc to search for perfect PIP boxes in the *A. citrulli* M6 genome, using the consensus sequence TTCGB-N15-TTCGB. Based on Koebnik *et al*. [32], we considered only those cases for which the distance between the end of the PIP box and the putative start codon was shorter than 650 nucleotides. This screen revealed a total of 78 PIP boxes (S4 Table), of which 41 correlated with significant regulation by HrpX (Table 2 and S4 Table). We used the PIP boxes of the aforementioned 41 genes/operons to determine the consensus PIP box of *A. citrulli* using the MEME suite (Fig 4).

**Fig. 4.**
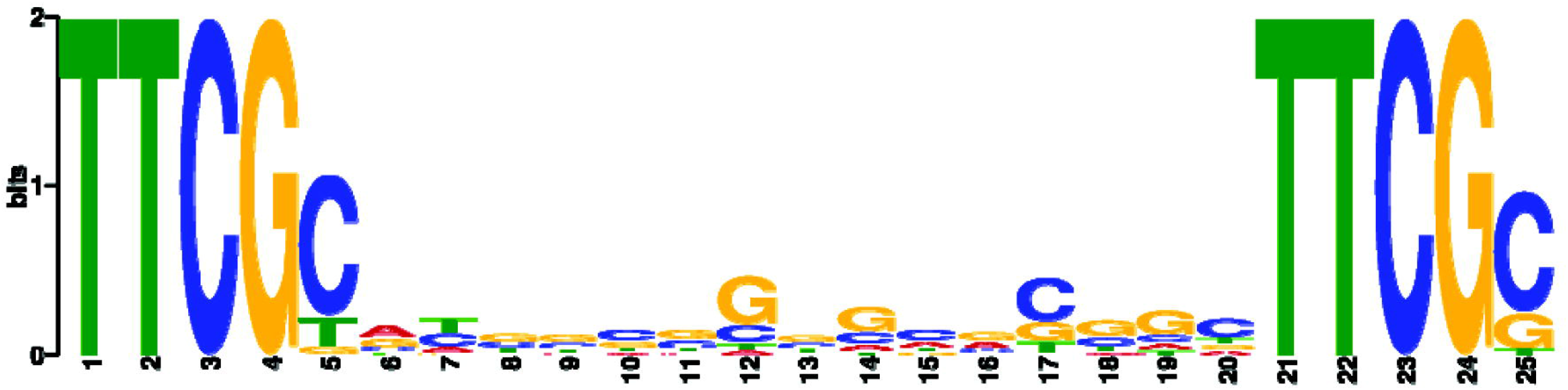
Sequence logo of the *Acidovorax citrulli* M6 plant-inducible promoter (PIP) box motif. The logo was generated with MEME-ChiP based on multiple alignment of the 41 perfect PIP boxes that were found to be associated with HrpX-regulated genes by RNA-Seq (see Table 2).

**Table 2.**
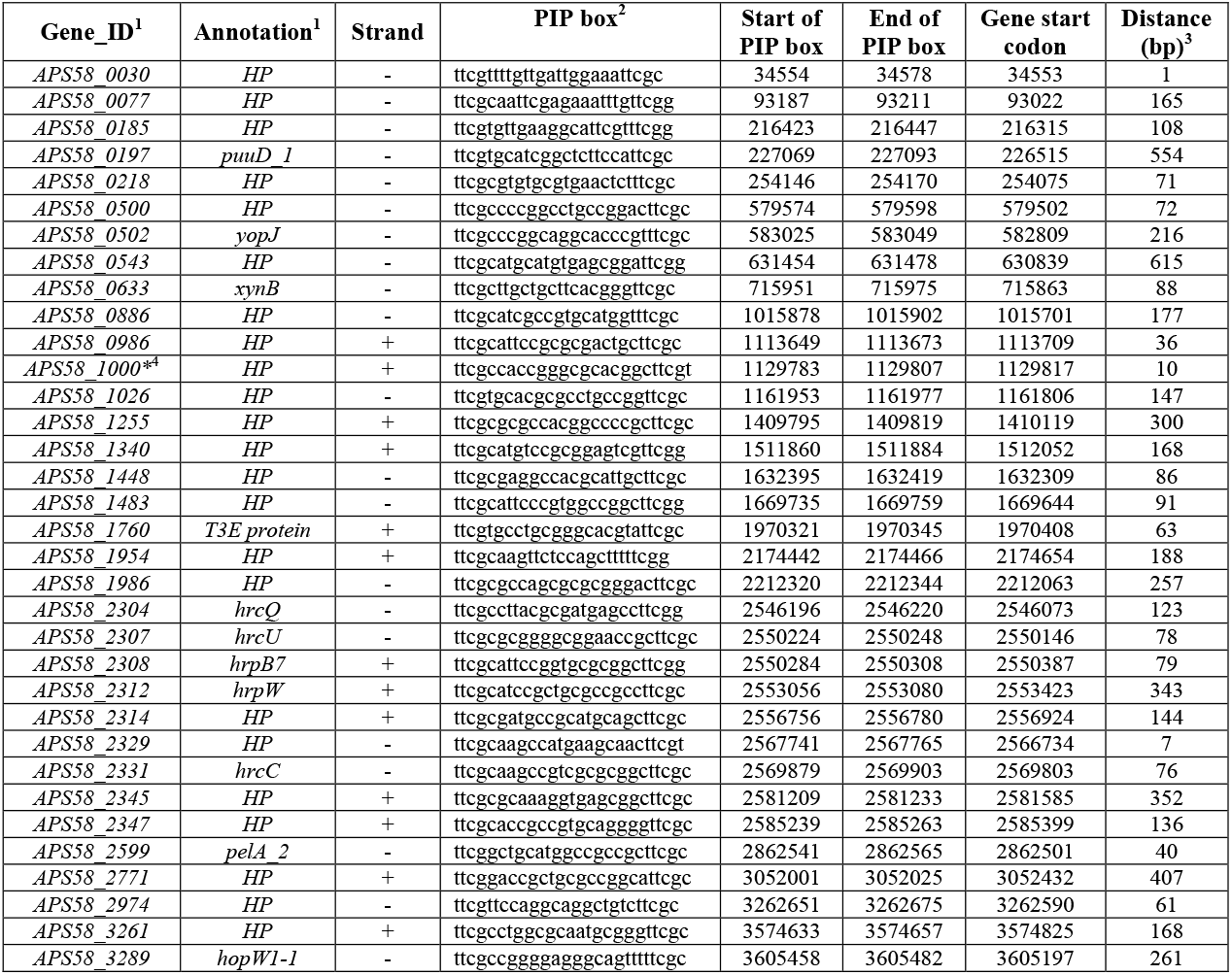
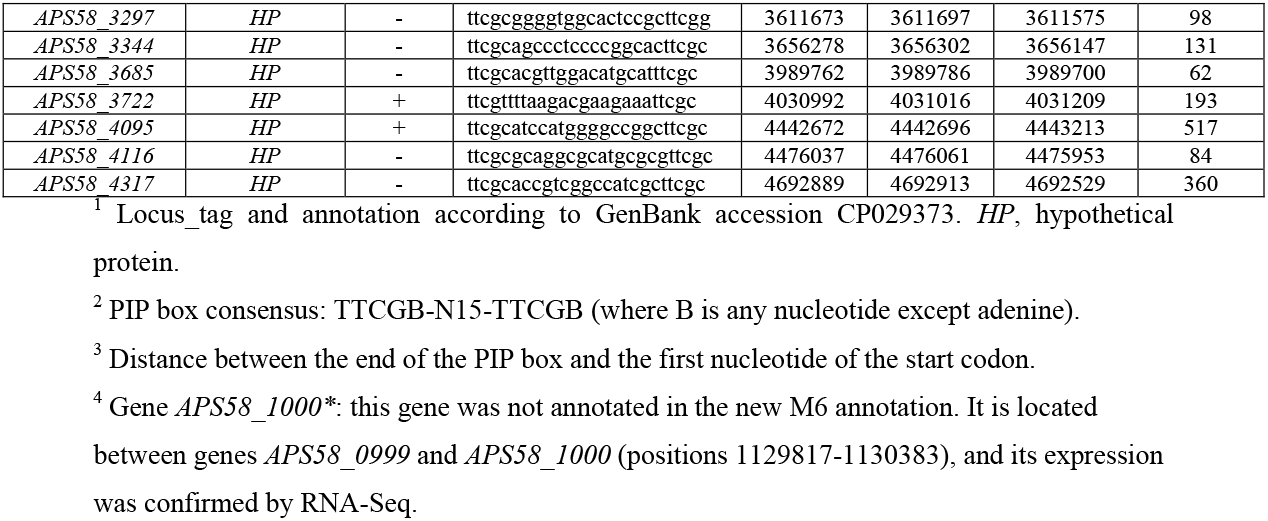
Perfect plant-inducible promoter (PIP) box sequences in genes that were shown to be regulated by HrpX in *Acidovorax citrulli* M6.

Importantly, some of the PIP boxes are upstream of operons, thus probably regulating the expression of more than one gene. We detected additional 25 genes [marked as (+) in the PIP box column of S3B Table] that are likely in PIP box-containing operons and showing higher expression in the wild-type strain relative to the *hrpX* mutant. It is also worth mentioning that eleven additional genes (some of which encoding T3Es) carrying PIP boxes showed higher expression values in the wild-type relative to the *hrpX* mutant in the RNA-Seq experiment, but were slightly below the level of statistical significance (S3A and S4 Tables).

### Establishment of a translocation assay for validation of *A. citrulli* T3Es

A critical prerequisite for the discovery of new T3Es is the availability of a suitable translocation assay. We assessed the possibility of exploiting the *avrBs2*-*Bs2* gene-for-gene interaction to test translocation of predicted *Acidovorax* T3Es into plant cells. The *X. euvesicatoria* AvrBs2 effector elicits an HR in pepper plants carrying the *Bs2* resistance gene [33]. A truncated form of this effector, carrying amino acids 62-574 (AvrBs2_62-574_), lacks the N-terminal T3S translocation signal, but retains the ability to elicit the HR when expressed in *Bs2* pepper cells [34]. The *avrBs2-Bs2* translocation assay is thus based on generation of plasmids carrying the candidate T3E (CT3E) genes fused upstream and in frame to the AvrBs2_62-574_. The plasmid is then mobilized into a *X. euvesicatoria* 85-10 *hrpG*ΔavrBs2* strain, that constitutively expresses *hrpG* and lacks *avrBs2*. The resulting strain is used to inoculate leaves of the pepper line ECW20R that carries the *Bs2* gene. If the AvrBs2_62-574_ domain is fused with a T3E gene, this elicits a Bs2-dependent HR [34]. Teper *et al*. recently used this reporter system to validate novel T3Es of the *X. euvesicatoria* strain 85-10 [28].

Given the close similarity between the T3SSs of *A. citrulli* and *Xanthomonas* spp., we hypothesized that the *X. euvesicatoria* T3S apparatus would recognize and translocate *A. citrulli* T3Es, and therefore, that the *avrBs2-Bs2* reporter system would be suitable for validating *A. citrulli* CT3E genes. To assess this hypothesis, we tested translocation of eight T3Es of *A. citrulli* showing similarity to known T3Es of other plant pathogenic bacteria. All tested fusions were translocated into pepper cells in a T3S-dependent manner and induced a Bs2-dependent HR in ECW20R pepper leaves. In contrast, HR was not detected when the fusions were tested in ECW30 leaves (lacking the *Bs2* gene), and when a *X. euvesicatoria hrpF* mutant (impaired in T3S) was used in these assays (Fig 5A). Overall, these results demonstrated the suitability of the *avrBs2-Bs2* assay for validation of *A. citrulli* CT3Es.

**Fig. 5.**
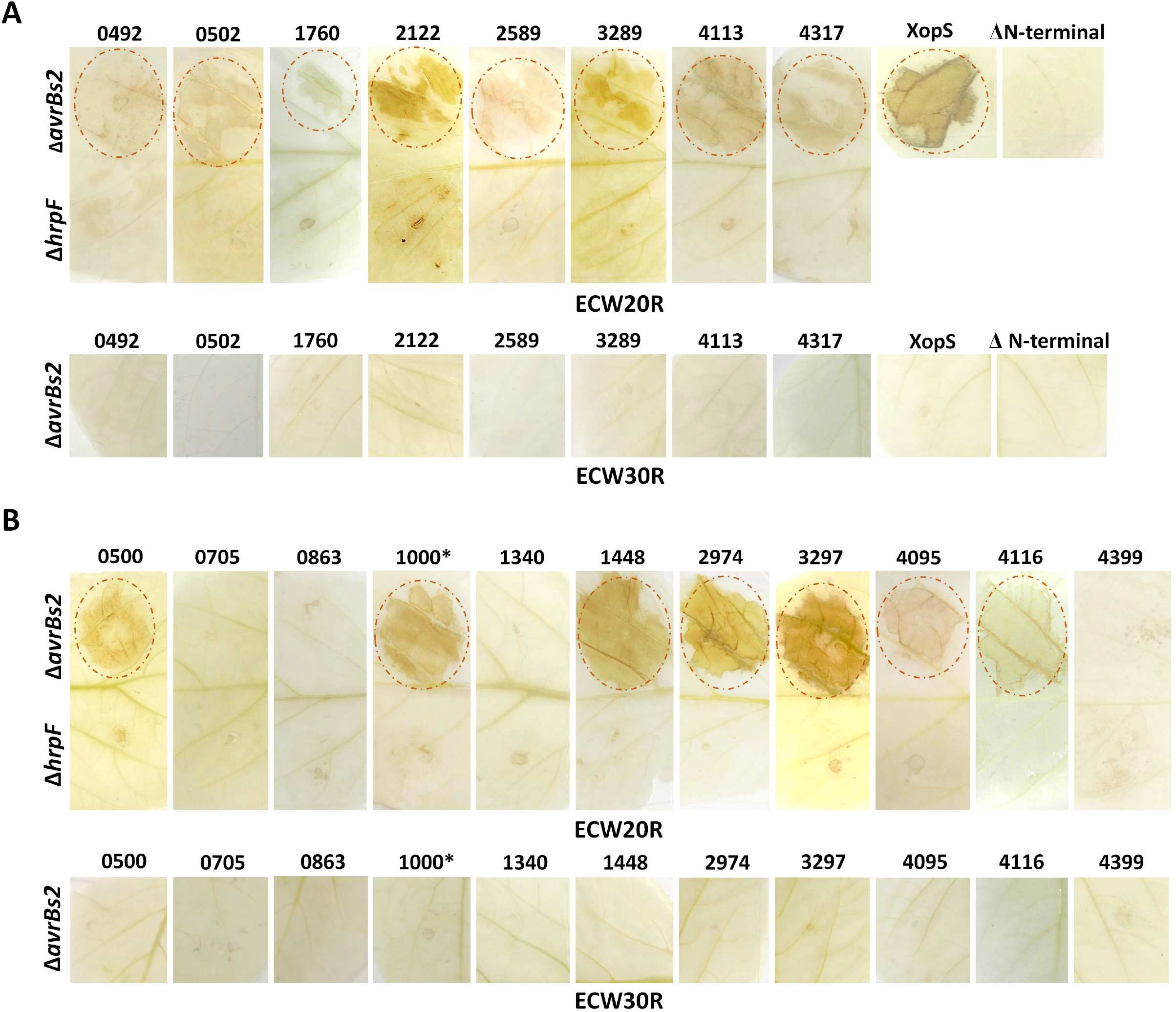
Translocation assays of T3Es of *Acidovorax citrulli* M6. (A) Selected T3Es based on sequence similarity to T3Es from other plant pathogenic bacterial species (see Table 1). (B) Candidate T3Es (CT3Es) selected from ML and RNA-Seq analyses. T3E/CT3E ORFs were cloned in plasmid pBBR1MCS-2 upstream to the AvrBs2_62-574_ domain, which elicits HR in ECW20R pepper plants carrying the *Bs2* gene, but not in ECW30R pepper plants that lack this gene. The plasmids were transformed into *Xanthomonas euvesicatoria 85-10-hrpG*-ΔavrBs2*, and the resulting strains were used to inoculate pepper plants. All known T3Es (A) and seven among eleven tested CT3Es (B) elicited HR in ECW20R but ECW30R leaves, similarly to the positive control XopS-AvrBs2_62-574_. Infiltrated areas are surrounded by red circles. No HR was induced when leaves were inoculated with a *X. euvesicatoria* mutant impaired in T3S *(ΔhrpF)* expressing T3E/CT3E-AvrBs2_62-574_ fusions. Also, no HR was induced following inoculation with *X. euvesicatoria 85-10-hrpG*-ΔavrBs2* without any plasmid (not shown) or with a plasmid expressing the AvrBs2_62-574_ domain alone (ΔN-terminal). Numbers at the top correspond to the locus_tag in strain M6 (for example, 0492 is gene *APS58_0492*).

### Seven novel T3Es of *A. citrulli* are translocated into plant cells

Following validation of the *avrBs2-Bs2* reporter assay for *A. citrulli* T3Es, we selected seven CT3Es based on results from the first ML run and RNA-Seq analysis. Four genes that were ranked relatively low in the ML were also included in these experiments to evaluate the quality of the ML prediction (Table 3 and S1 Table). All seven CT3E genes, but not the low-ranked ML genes, were translocated (Fig 5B). The validated genes were annotated as hypothetical proteins, had a predicted PIP box, were shown to be positively regulated by HrpX, and ranked high in the ML run (Table 3 and S1 Table). Importantly, the gene *APS58_1340*, which contains a PIP box in its promoter region and its expression is regulated by HrpX (Table 3) was not translocated, indicating that these two parameters alone are not sufficient for accurate prediction of T3Es.

**Table 3.** Candidate T3E genes of *Acidovorax citrulli* M6 that were tested in the *avrBs2-Bs2* translocation assays.

| Gene ID<sup>1</sup> | Product | ML<sup>2</sup> | PIP<sup>3</sup> | RSEQ<sup>4</sup> | TRA<sup>5</sup> |
| --- | --- | --- | --- | --- | --- |
| *APS58_0500* | Hypothetical protein<sup>7</sup> | 39/48 | + | + | + |
| *APS58_0705* | GrxD, glutaredoxin-4 | 91/2203 | - | - | - |
| *APS58_0863* | Hypothetical protein | 64/749 | - | - | - |
| *APS58_1000*<sup>*6</sup> | Hypothetical protein<sup>7</sup> | 21/* | + | + | + |
| *APS58_1340* | Hypothetical protein | 84/535 | + | + | - |
| *APS58_1448* | Hypothetical protein<sup>7</sup> | 17/104 | + | + | + |
| *APS58_2974* | Hypothetical protein<sup>7</sup> | 19/29 | + | + | + |
| *APS58_3297* | Hypothetical protein<sup>7</sup> | 20/61 | + | + | + |
| *APS58_4095* | Hypothetical protein<sup>7</sup> | 31/46 | + | + | + |
| *APS58_4116* | Hypothetical protein<sup>7</sup> | 11/43 | + | + | + |
| *APS58_4399* | Hypothetical protein | 174/739 | - | - | - |

<sup>1</sup> Gene IDs are according to the annotation of the *A. citrulli* M6 chromosome (GenBank accession CP029373).
<sup>2</sup> ML: rankings in first/second machine learning (ML) runs. \*, gene *APS58\_1000\** was not included in the second ML run (see below).
<sup>3</sup> PIP: presence (+) or absence (-) of perfect plant-inducible promoter (PIP) box in the promoter region.
<sup>4</sup> RSEQ: significantly reduced expression in the *hrpX* mutant relative to the wild type (+)/no significant differences between strains (-).
<sup>5</sup> TRA: translocated (+)/non-translocated (-) in *avrBs2-Bs2* translocation assays (rows of validated genes are shaded with gray).
<sup>6</sup> *APS58\_1000\**: this gene ranked high in the first ML but was not annotated in the recent annotation of *A. citrulli* M6, although its expression was confirmed by RNA-Seq. Its ORF is located between genes *APS58\_0999* and *APS58\_1000* (positions 1129817-1130383).
<sup>7</sup> These genes were detected only in plant pathogenic *Acidovorax* species.

BlastP analyses of the seven newly identified T3E genes revealed strong similarity only to hypothetical proteins of plant pathogenic *Acidovorax* species. The fact that no homologs for these genes were detected in non-pathogenic *Acidovorax* strains (despite the availability of more than 70 genomes of such spp.) or in other plant pathogenic bacterial species suggests a specific and unique role for their products in *Acidovorax* pathogenicity. These seven genes were detected also in AAC00-1 (S2 Table) and in all other group I and II genomes available in NCBI. Some of them were widely distributed among other plant pathogenic *Acidovorax* species. For instance, *APS58_4095* was also detected in *A. oryzae* and in *A. cattleyae*, and homologs with less than 60% query coverage were also present in *A. konjaci, A. anthurii* and *A. valerianellae*. In contrast, *APS58 2974* was not detected in *Acidovorax* spp., other than *A. citrulli* and *A. avenae* (S2 Table). Searches for conserved domains in these T3Es did not provide any insight.

### Assessment of localization of three of the newly identified T3Es

We attempted to assess the subcellular localization of three of the newly identified T3Es, APS58_0500, APS58_1448 and APS58_4116. Prediction of subcellular localization using the Plant-mPLoc server indicated that the three effectors could localize to the nucleus. Browsing these T3Es with the LogSigDB server revealed endoplasmic reticulum (ER) localization signals in the three effectors, and nuclear localization signals in APS58_0500 and APS58_4116.

We assessed localization of these effectors fused to the yellow fluorescent protein (YFP) in *Nicotiana benthamiana* leaves following transient expression by agroinfiltration. Based on the aforementioned predictions, in first experiments the leaves were co-infiltrated with *A. tumefaciens* carrying the ER marker mRFP-HDEL, and were also stained with DAPI for nucleus localization. Representative images from these experiments are shown in Fig 6. The results suggested that the three effectors could interact with the ER, but only APS58_0500 and APS58_1448 partially localized to the nucleus, including in clearly visible nuclear foci (Fig 6).

**Fig. 6.**
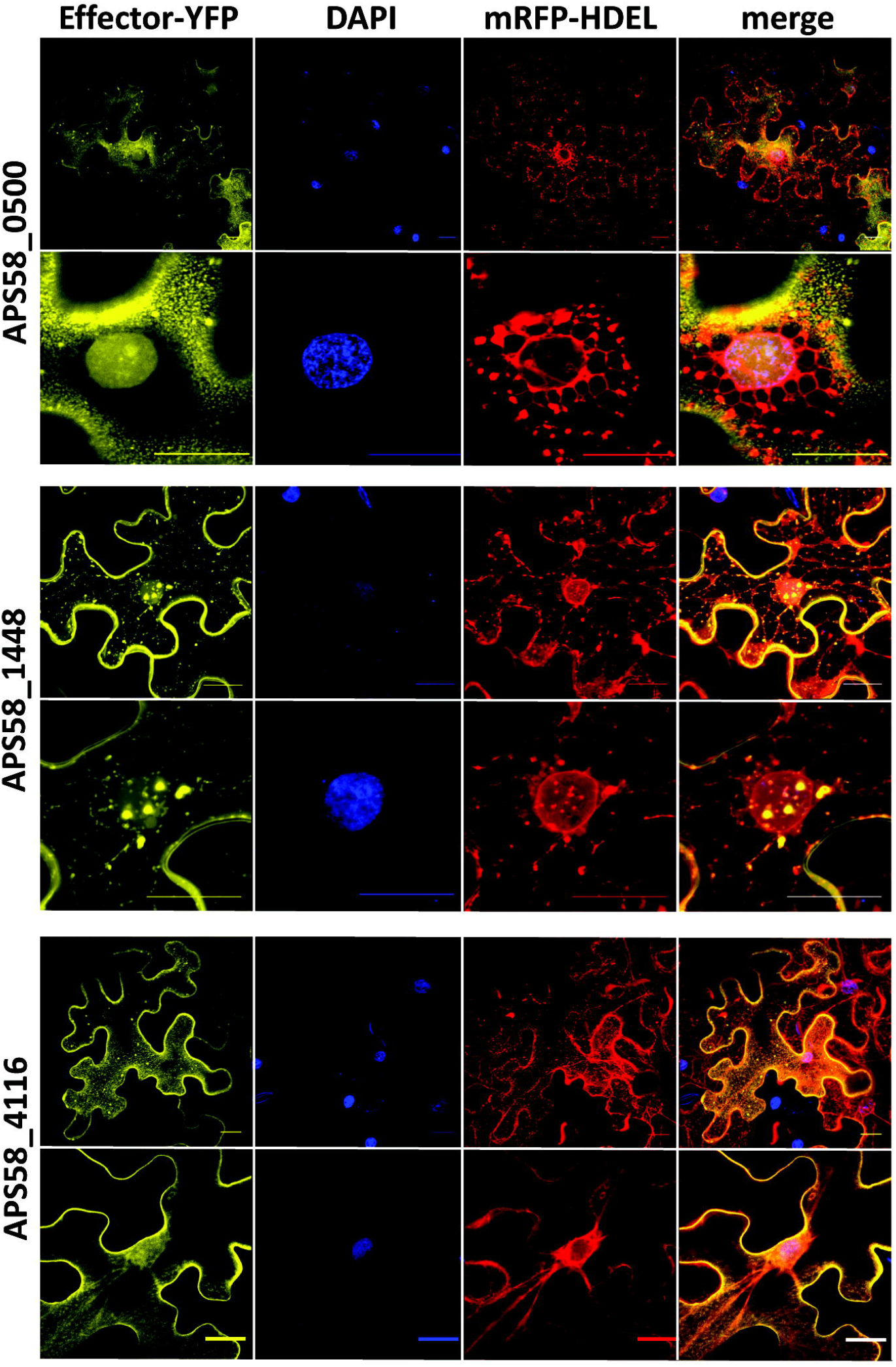
Transient expression of *Acidovorax citrulli* T3Es in *Nicotiana benthamiana*. The T3E genes *APS58_0500, APS58_4116* and *APS58_1448*, identified by ML and RNA-Seq and validated in translocation assays, were cloned in the binary vector pEarleyGate101, fused to the C-terminus of YFP. The plasmids were transformed into *Agrobacterium tumefaciens* GV3101, and the resulting strains were used for transient expression in *N. benthamiana*. Leaves were co-inoculated with *A. tumefaciens* GV3101 carrying the mRFP-HDEL endoplasmic reticulum marker and stained with DAPI for visualization of plant cell nuclei. Samples were visualized in a Leica SPE confocal microscope 48 h after inoculation. Bars at the right bottom of each picture, 20 μm.

In a second set of experiments, the YFP-fused effectors were co-infiltrated with free-mCherry, localized mainly in the cytosol and in the nucleus, HDEL-mCherry, localized to the ER, and the membrane-bound protein SlDRP2A (L. Pizarro and M. Bar, unpublished results). Representative images from these experiments are shown in S2-S4 Figs for APS58_0500, APS58_1448 and APS58_4116, respectively. The three effectors partially co-localized with the membrane-bound protein SIDRPA, as evidenced by the Pearson correlation coefficients (0.40±0.024 for APS5_0500, 0.49±0.040 for APS58_1448, and 0.53±0.037 for APS58_4116). Since APS58_0500 appeared to have a stronger membrane localization, we used the classical plasma membrane microdomain protein Flot1 [35] as an additional membrane control marker. Indeed, APS58_0500 had an expression pattern that was highly similar to that of Flot1 (compare top and bottom panels in S2 Fig). In agreement with the first set of experiments (Fig 6), APS58_0500 (S2 Fig) and APS58_1448 (S3 Fig) partially localized to the nucleus. On the other hand, these experiments confirmed that only APS58_4116 partially interacted with the ER, mostly at the nuclear envelope (Fig 6 and S4 Fig; Pearson coefficient with HDEL-mCherry 0.53±0.037). None of the effectors was shown to have a significant cytosolic presence: the Pearson coefficient with mCherry was lower than 0.12 for APS_0500 and APS_4116, while for APS_1448 the coefficient was 0.53±0.037, due to the strong nuclear presence of this effector, as indicated above. Overall, we can conclude that the three effectors are associated with the plasma membrane, APS58_0500 and APS58_1448 partially localize to the nucleus, and APS_4116 partially interacts with the ER.

### Generating an improved list of candidate T3Es of *A. citrulli* M6 with a second ML run

Machine learning can be improved after refinement of features specific to the studied pathogen. Thus, we carried out a second ML run using the *A. citrulli* M6 genome. The main differences between the first and second ML runs were: (i) the second run was done on the M6 genome [9], which by this time was fully assembled; (ii) we included the seven novel T3Es identified in this study in the positive set and the four ORFs that were found not to be translocated were added to the negative set; (iii) in the positive set we included ORFs with high sequence similarity to known effectors from other bacteria, based on our homology search results (Table 1); and (iv) we used HrpX-mediated regulation as an additional feature used to train the classifier. The results are summarized in S5 Table. Most known/validated T3Es ranked among the top 100 hits, and among the top 40 hits, 34 were known/validated T3Es. Importantly, some genes with high propensity to encode T3Es (ranking among the top 60 in the second ML run) did not appear among the top 200 hits in the first ML list (Table 1 and S1 Table), thus supporting the higher reliability of the new list relative to the first prediction.

Among the top 100 hits of the second ML run, there were 37 genes that matched to hypothetical proteins from the public database, with no similarity evidence to suggest a T3E nature. Since this was the case of the seven T3Es validated in this study, it is possible that some of these genes encode previously undiscovered T3Es. In this regard, it is worth mentioning genes *APS58_1954, APS58_1986, APS58_3685, APS58_0987* and *APS58_1694* (ranking at positions 20, 27, 57, 62 and 83 in the second ML, respectively). While *APS58_1694* shares similarity only with hypothetical proteins of plant pathogenic *Acidovorax* species, the first four also share similarities to hypothetical proteins of other plant pathogenic genera (eg., *Xanthomonas, Ralstonia, Pseudomonas* and/or *Erwinia*). These genes also showed increased expression in wild-type M6 relative to the *hrpX* mutant, and have PIP boxes in their promoter region. Therefore, these genes are strong candidates for further experimental validations.

## Discussion

Type III effectors (T3Es) play a dual role in the interaction between many Gram-negative plant pathogenic bacteria and plants: while they collectively promote virulence on susceptible plants, some may induce effector-triggered immunity (ETI) in plants carrying the corresponding resistance (*R*) genes. *R* genes provide resistance against economically important pathogens and have been mobilized to commercial crop varieties by breeding programs. Thus, this is one of the most important means of disease management [36, 37].

*Acidovorax citrulli* requires a functional type III secretion (T3S) system for pathogenicity [10]. The main objective of this study was to significantly advance the current knowledge about the arsenal of T3Es of *A. citrulli*. Among well-investigated plant pathogenic bacteria, the pools of T3Es vary from only few effectors in phytopathogenic bacteria from the *Enterobacteriaceae* family, to approximately 20 to 40 in strains of *P. syringae* and *Xanthomonas* spp. [21, 28, 40–44] and an average of over 75 in *R. solanacearum* isolates [45, 46]. Thus, we hypothesized that the repertoire of *A. citrulli* T3Es could be much larger than the eleven T3E genes identified in the group II strain AAC00-1 [27].

As a first approach to uncover the arsenal of *A. citrulli* T3Es, we used a genome-wide machine learning (ML) algorithm to determine the propensity of ORFs to encode T3Es. In parallel, we looked carefully at the annotation of the group I model strain of *A. citrulli*, M6, and carried out BlastP analyses of the genes encoding hypothetical proteins or functions that could infer effector activity. These analyses revealed 51 putative T3E genes that shared different levels of similarity with known effector genes from *Xanthomonas* spp., *R. solanacearum* and/or *P. syringae* strains (Table 1). Homologs for most of these T3E genes and for those identified in the present study were also detected in other plant pathogenic *Acidovorax* species (S2 Table).

To identify new T3E genes of *A. citrulli*, we also used RNA-Seq to identify HrpX-regulated genes. Based on the knowledge accumulated with *Xanthomonas* spp. and *R. solanacearum* [22, 32, 47, 48], we expected that most genes encoding T3SS components and some T3Es of *A. citrulli* would be under the direct regulation of HrpX. This assumption was strengthened in preliminary experiments comparing gene expression between a wild-type and a *hrpX* mutant strain (Fig 1C). As previously mentioned, Zhang *et al*. recently showed that HrpX controls the expression of one T3E gene in the group II strain, Aac5 [26].

The RNA-Seq approach revealed 159 genes showing significantly reduced expression in the *hrpX* mutant, while 28 genes had significantly increased expression in the mutant (S3 Table). These numbers are similar to those reported in gene expression studies carried out with *Xanthomonas* spp. HrpX and with *R. solanacearum* HrpB. For instance, microarray analyses of *Xanthomonas axonopodis* pv. *citri (Xac)* in XVM2 medium revealed that 181 genes were up-regulated by HrpX, while 5 to 55 genes (depending on the time point) were down-regulated by this transcriptional regulator [47]. Occhialini *et al*. found 143 HrpB up-regulated genes and 50 HrpB down-regulated genes in *R. solanacearum* [48]. In these, as well as in several other studies, HrpX/HrpB was found to regulate the expression of most genes encoding T3S components and accessory proteins as well as several T3E genes [21, 22, 49]. In line with this background, among the 159 HrpX up-regulated genes found in our study, 20 encoded *hrp/hrc/hpa* genes and 27 encoded T3E genes. Interestingly, *hrcC* was a member of the *A. citrulli* HrpX regulon. *hrcC* expression in *X. euvesicatoria* is directly regulated by HrpG, in an HrpX-independent manner [31]. In contrast, in *R. solanacearum, hrcC* is regulated by HrpX [49, 50] as we found in *A. citrulli* M6.

In *Xanthomonas* spp. and in *R. solanacearum*, the HrpX/HrpB regulon includes many genes that are not involved in T3S [21, 47, 49]. A similar picture emerged from our study, where HrpX was shown to regulate genes involved in transmembrane transport, including several ABC transporters and permeases as well as transcriptional regulators. Among the HrpX up-regulated genes we also detected several genes whose products are putatively secreted by type II secretion (T2S). These included genes encoding 1-4-β-xylanase *(xynB)*, pectate lyase and a protein with similarity to a family of S1 extracellular serine proteases (S3E Table). HrpX regulation of genes encoding type II-secreted enzymes was also demonstrated in *Xanthomonas* spp. and in *R. solanacearum* [22, 47, 51–54].

Among the 159 HrpX up-regulated genes *in A. citrulli*, more than 60 carried perfect PIP boxes in their promoter region or were part of operons carrying perfect PIP boxes (Table 2; S3 and S4 Tables). Although some other genes may carry imperfect PIP boxes and may be directly regulated by HrpX, this result suggests that many of the HrpX up-regulated genes are indirectly regulated by this transcriptional factor. This is a reasonable assumption, considering that among the genes that are up- and down-regulated by HrpX, there are several transcriptional regulators. For instance, genes encoding transcriptional factors belonging to the LysR *(APS58_0949* and *APS58_2039*), IclR *(APS58_1263)*, FmbD *(APS58_1340)* and TetR *(APS58_3638)* families were shown to be up-regulated by HrpX. In contrast, two genes encoding DNA-binding response regulators, homologous to PhoP *(APS58_0821)* and FixJ *(APS58_1682)* were HrpX-down-regulated (S2B Table).

After demonstrating the suitability of the *avrBs2-Bs2* T3E translocation assay with eight known T3Es, we used the data obtained from the first ML run and the RNA-Seq analysis to select seven *A. citrulli* M6 ORFs for experimental validation (Fig 5). We validated translocation of the seven candidates, thus demonstrating the strength of combining ML and RNA-Seq for identifying T3E genes. Importantly, the lack of translocation of the four ORFs that received relatively low scores in the first ML run strengthened the suitability of our combined computational/experimental approach.

Remarkably, the seven effectors identified in this study were up-regulated by HrpX and carried PIP boxes in their promoter regions, while among the four nonvalidated genes, only one had these traits (Table 3). An interesting trait of the seven new T3Es was that they share significant similarity only with hypothetical proteins of other plant pathogenic *Acidovorax* strains (Table 3 and S2 Table). This strongly supports that these effectors are unique to plant pathogenic *Acidovorax*. Importantly, a second ML run, informed by the knowledge accumulated from this study, revealed additional genes that were ranked in relatively high positions and encoded hypothetical proteins that occur only in plant pathogenic *Acidovorax* or in other plant pathogenic bacteria (S5 Table). These represent high priority CT3Es for future experimental validation assays. This emphasizes one benefit of the ML approach: its ability to integrate novel knowledge in the prediction algorithm.

Another interesting characteristic of the new T3Es discovered in this study is their relatively small size. Based on the annotation of the M6 genome, the average and median lengths of *A. citrulli* M6 T3Es are 387.7 and 345 amino acids (a.a.), respectively. Except for *APS 4116* that encodes a 347-a.a. protein, the size of the six other new T3Es ranged from 113 a.a. (APS58_4095) to 233 a.a. (APS58_0500) (S5 Fig). In the public database (GenBank), there are several examples of small T3Es from plant pathogenic bacteria, including *Xanthomonas* AvrXv3 (most having 119 a.a.), *Pseudomonas syringae* HopAF1 (112-291 a.a.), HopBF1 (125-207 a.a.), HopF2 (177-280 a.a.), HopH1 (201-218 a.a.) and AvrRpt2 (222-255 a.a.), and the *R. solanacearum/Xanthomonas* HopH1 homologs (155-218 a.a.).

In this study we assessed plant cell localization of three of the new T3Es validated in translocation assays, APS58_0500, APS58_1448 and APS58_4116. Utilization of subcellular localization prediction tools and confocal microscopy of *N. benthamiana* agro-infiltrated leaves strongly suggest that the three tested effectors interact with the plasma membrane (S2-S4 Figs), with APS58_0500 remarkably mimicking the localization of the classical non-clathrin mediated endocytic system protein, Flot1 [35]. While APS58_4116 interacted with the endoplasmic reticulum (Fig 6 and S2 Fig), effectors APS58_0500 and APS58_1448 partially localized to the nucleus (Fig 6; S3 and S4 Figs). Interestingly, and in line with the predicted nuclear localization of these effectors, BlastP showed that APS58_0500 has low similarity with an ATP-dependent RNA helicase of the Metazoa organism *Clonorchis sinensis* (query cover, 38%; e-value, 0.23), while APS58_1448 has low similarity to a transcriptional regulator of the bacterium *Hoeflea halophila* (query cover, 54%; e-value, 0.19-0.53), suggesting possible functions the *Acidovorax* effectors might execute upon entrance into the plant cell nucleus.

In conclusion, we have combined sequence analysis, ML and RNA-Seq approaches to uncover the arsenal of T3Es of the group I model strain of *A. citrulli*, M6, including discovery of new T3Es that appear to be unique to plant pathogenic *Acidovorax* spp. Further characterization of the novel T3Es identified in this study may uncover new host targets of pathogen effectors and new mechanisms by which pathogenic bacteria manipulate their hosts. We also demonstrated the suitability of a translocation reporter system for validation of *A. citrulli* T3Es, which we expect, will be very helpful to the *Acidovorax* research community. Until recently it was assumed that

*A. citrulli* strains (and in general plant pathogenic *Acidovorax* strains) possess little over ten T3E genes. However, from this study it is clear that the *A. citrulli* pan-genome encodes more than 50-60 T3Es. Therefore, the *A. citrulli* T3E repertoire is larger than those of most well-characterized plant pathogenic bacteria, including plant pathogenic *Enterobacteria, P. syringae* pathovars, *Xanthomonas* spp. and, and closer in numbers to the T3E repertoires of *R. solanacearum*. Moreover, the second ML run suggested that *A. citrulli* may possess yet unrevealed T3E genes. Importantly, among the 58 known T3E genes of *A. citrulli* M6, ten (17.2%) appear to be unique to group I strains. On the other hand, the group II model strain, AAC00-1, carries T3E genes that are absent or nonfunctional in group I strains as shown in our previous report [27] and in this study (Table 1 and S1 Table). Thus, it is logical to assume that the variability in T3E content between group I and II strains plays a critical role in shaping the differences in host-preferential association between the groups. Despite this, more research is needed to test this hypothesis, and to understand the mode of action and contribution of individual effectors to the virulence of *A. citrulli*.

## Methods

### Bacterial strains and plasmids

Bacterial strains and plasmids used in this study are listed in S6 Table. Unless stated otherwise, *Acidovorax citrulli* strains were grown at 28 °C in nutrient broth (NB; Difco Laboratories, Detroit, Michigan) or nutrient agar (NA; NB containing 15 g/L agar). For RT-PCR, qRT-PCR and RNA-seq experiments, *A. citrulli* strains were grown in XVM2 medium [31]. *Xanthomonas euvesicatoria, Agrobacterium tumefaciens* and *Escherichia coli* strains were cultured on Luria-Bertani (LB) medium [55] at 28 °C for *X. euvesicatoria* and *A. tumefaciens*, and 37 °C for *E. coli*. When required, media were supplemented with the following antibiotics: ampicillin (Ap, 100 μg/mL for *E. coli* and 200 μg/mL for the others), rifampicin (Rif, 50 μg/mL), kanamycin (Km, 50 μg/mL), and gentamycin (Gm, 50 μg/mL for *A. citrulli* and 10 μg/mL for the others).

### Molecular manipulations

Routine molecular manipulations and cloning procedures were carried out as described [55]. T4 DNA ligase and restriction enzymes were purchased from Fermentas (Burlington, Canada). AccuPrep^®^ Plasmid Mini Extraction Kit and AccuPrep^®^ PCR Purification Kit were used for plasmid and PCR product extraction and purification, respectively (Bioneer Corporation, Daejeon, Republic of Korea). DNA was extracted with the GeneElute bacterial genomic DNA Kit (Sigma-Aldrich, St. Louis, MO, USA). PCR primers were purchased from Sigma-Aldrich and are listed in S7 Table. PCR reactions were performed with the Readymix Red Taq PCR reactive mix (Sigma-Aldrich) or with the Phusion high-fidelity DNA polymerase (Fermentas, Waltham, MA, USA) using an Eppendorf (Hamburg, Germany) thermal cycler. Sequencing of PCR fragments and constructs was performed at Hy Laboratories (Rehovot, Israel). *Escherichia coli* S17-1 λpir, DH5-α and DB3.1 strains were transformed using an Eppendorf 2510 electroporator according to manufacturer’s instructions. Plasmid mobilizations to *A. citrulli* and *X. euvesicatoria* strains were done by bi-parental mating as described [56]. *A. tumefaciens* cells were transformed by the heat shock method [57].

### Machine learning classifications

In order to predict T3Es, we applied ML classification algorithms, which are similar to the ones we have previously described [28, 29, 58, 59]. The first ML run was used to search for T3Es in the AAC00-1 genome (GenBank accession CP000512.1).

The training data included 12 ORFs that were known as T3Es (see in Results). The negative set included 2,680 ORFs that had high similarity (E-value less than 0.001) to ORFs in the non-pathogenic *E. coli* K12 genome (accession number NC_000913.3). The positive and negative ORFs are marked in S1 Table. For this ML, 71 features were used, including homology (to known effectors or to bacteria without T3SS), composition (amino acid composition, GC content), location in the genome (e.g., distance from known T3Es), and the presence of a PIP box in the promoter region. The complete list of features is given in S8 Table. Features were extracted using in-house Python scripts. The outcome of the ML run is a score for each ORF, reflecting its likelihood to encode a T3E. We evaluated several classification algorithms: random forest [60], naïve Bayes [61], support vector machine (SVM; [62]), K nearest neighbors (KNN), linear discriminate analysis (LDA), logistic regression (all three described in Hastie *et al*. [63]), and Voting, which aims to predict averaging over all other ML algorithms. For each run, feature selection was performed. The ML algorithms and feature selection were based on the Scikit-learn module in Python [64]. The area under the curve (AUC) score over 10-fold cross-validation was used as a measure of the classifier performance. The first ML run was based on the random forest classifier which gave the highest AUC (0.965).

The second ML run was similar to the first, with the following modifications. First, the classifiers were run on the M6 genome (GenBank accession CP029373). Second, the positive set included ORFs that were validated as T3Es in this study and ORFs with high sequence similarity to known effectors as described in Table 1. The negative set included 2,570 ORFs (S5 Table). The four ORFs that were experimentally shown not to be T3Es were also included in this negative set. Third, the expression data from the RNA-Seq analysis (HrpX regulation) were added as a feature. Fourth, the PIP box feature was updated to reflect the PIP box as inferred from promoter regions of *Acidovorax* T3Es (see additional bioinformatics tools below). The second ML run was based on Voting classifier, which included all the classifiers specified above, as it gave the highest AUC among all the classifiers. The AUC for this second ML run was 0.999.

### Generation of *A. citrulli* mutants and complemented strains

*Acidovorax citrulli* M6 mutants disrupted in *hrpX (APS58_2298)* and *hrpG (APS58_2299)* genes were generated by single insertional mutagenesis following single homologous recombination. Internal fragments of the *hrpX* (383 bp) and *hrpG* (438 bp) ORFs carrying nucleotide substitutions that encode early stop codons, were PCR-amplified and inserted into the *Bam*HI/EcoRI site of the suicide plasmid pJP5603 [65]. The resulting constructs were transformed into *E. coli* S17-1 λpir, verified by sequencing, and mobilized into *A. citrulli* M6 by bi-parental mating. Transconjugants were selected by Km selection. Disruption of the target genes by single homologous recombination and plasmid insertion was confirmed by PCR and sequencing of amplified fragments. To generate complemented strains for mutants disrupted in *hrpX* and *hrpG* genes, the full ORFs of these genes (1407 pb and 801 bp, respectively) were PCR-amplified and cloned into the EcoRI/*Bam*HI sites of pBBR1MCS-5 [66]. The generated plasmids were transformed into *E. coli* S17-1 λpir, verified by sequencing, and transferred by bi-parental mating into the corresponding M6 mutant strains. Complemented strains were selected by Gm resistance and validated by PCR.

### Infiltration of melon and pepper leaves with *A. citrulli* strains

Melon *(Cucumis melo)* cv. HA61428 (Hazera Genetics, Berurim, Israel) plants were grown in a greenhouse at ~28 °C. Pepper *(Capsicum annum)* cv. ECW20R and ECW30 [67] plants were grown in a growth chamber (16 h/26 °C in the light; 8h/18°C in the dark; relative humidity set to 70%). The three youngest, fully expanded leaves of 3-week-old melon and 5-week-old pepper plants were syringe-infiltrated in the abaxial side with bacterial suspensions of *A. citrulli* strains containing 10^8^ colony forming units (cfu)/mL in 10 mM MgCl2. Phenotypes were recorded 3 and 4 days after inoculation (d.a.i.), for melon and pepper leaves, respectively. For a better visualization of HR symptoms in pepper leaves, the infiltrated leaves were soaked in an acetic acid:glycerol:water solution (1:1:1 v/v) for 4 h and then transferred to ethanol and boiled for 10 min. Experiments were repeated twice with similar results.

### RNA isolation, cDNA synthesis and reverse transcription-PCR (RT-PCR)

*Acidovorax citrulli* M6 and *hrpX* mutant were grown at 28 °C in 5 mL of XVM2 medium for 72 h. Total RNA was isolated using TRI reagent (Sigma-Aldrich) and Direct-zol RNA miniprep kit (Zymo Research, Irvine, CA, USA) according to manufacturer’s instructions. Samples were treated with RNase free DNase using Turbo DNA-free kit (Invitrogen, Carlsbad, CA, USA). RNA concentration was quantified using a Nanodrop DS-11 FX (Denovix, Wilmington, Delaware) and RNA integrity was assayed on 1% agarose gels. RNA was reverse transcribed into cDNA using a High Capacity cDNA Reverse Transcription Kit (Applied Biosystems). Semiquantitative RT-PCR analysis was performed using 1 μg of cDNA or gDNA (as positive control for amplification), 0.6 pmol of selected primer, the Phusion High-Fidelity DNA Polymerase (ThermoFisher Scientific, Waltham, MA, USA), and the following conditions: 98 °C for 15 min, followed by 35 cycles of 98 °C for 30 s, 60 °C for 30 s and 72 °C for 15 s. The *A. citrulli GADPH* housekeeping gene [68] was used as reference. The relative amount of amplified DNA was assayed on 2% agarose gels.

### RNA-Seq and quality analysis

Total RNA of wild-type M6 and *hrpX* mutant strains was isolated as described above for RT-PCR experiments. Three independent RNA extractions were obtained for each strain. Ribosomal RNA was depleted using the MICROB Express Bacterial mRNA Purification kit (Ambion, Foster City, CA, USA). The integrity and quality of the ribosomal depleted RNA was checked by an Agilent 2100 Bioanalyzer chip-based capillary electrophoresis machine (Agilent Technologies, Santa Clara, CA, USA). RNA sequencing was carried out at the Center for Genomic Technologies at The Hebrew University of Jerusalem (Jerusalem, Israel). The samples were used to generate whole transcriptome libraries using the NextSeq 500 high output kit (Illumina, San Diego, CA, USA) with a NextSeq 2000 sequencing instrument (Illumina). The cDNA libraries were quantified with a Qubit 2.0 Fluorometer (Invitrogen) and their quality was assessed with an Agilent 2200 TapeStation system (Agilent Technologies). One of the *hrpX* mutant libraries was removed from further analysis due to low quality. Raw reads (fastq files) were further inspected with FastQC v0.11.4 [69]. They were trimmed for quality and adaptor removal using Trim Galore default settings: trimming mode, single-end; Trim Galore version 0.4.3; Cutadapt version 1.12; Quality Phred score cutoff, 20; quality encoding type selected, ASCII+33; adapter sequence, AGATCGGAAGAGC (Illumina TruSeq, Sanger iPCR; auto-detected); maximum trimming error rate; 0.1; minimum required adapter overlap (stringency), 1 bp. An average of 0.6% of the reads were quality trimmed and 57% of the reads were treated for adaptor removal.

### Mapping of RNA-Seq reads on the *A. citrulli* M6 genome and differential expression analysis

Cleaned reads (~20 million per sample) were mapped against the latest version of the *A. citrulli* M6 genome (CP029373) using STAR v 2.201 [70]. Mapping files were further processed for visualization by Samtools Utilities v 0.1.19 [71]. The resulting Bam files were used to improve gene and operon predictions along the genome using cufflinks v2.2.1 followed by cuffmerge without a guiding reference file [72]. Uniquely mapped reads per gene were counted twice [once using the original submitted annotation file (orig.gff), and then using the merged annotations by cufflinks-cuffmerge (merged.gff)] using HTSeq-count [73]. Differential expression analysis was performed using the DESeq2 R package [73]. Differentially expressed genes were defined as those genes with a fold-change higher than 2, and a *P* value lower than 0.05.

### Validation of RNA-Seq results by quantitative real-time PCR (qRT-PCR)

RNA-seq data were verified by qRT-PCR using specific primers of selected genes (S7 Table). Bacterial growth, RNA isolation and cDNA synthesis were as described above for RT-PCR and RNA-Seq experiments. qRT-PCR reactions were performed in a Light Cycler 480 II (Roche, Basel, Switzerland) using 1 μg of cDNA, 0. 6 pmol of each primer and the HOT FIREPol EvaGreen qPCR Mix Plus (Solis BioDyne, Tartu, Estonia), and the following conditions: 95 °C for 15 min (1 cycle); 95 °C for 15 s, 60 °C for 20 s and 72 °C for 20 s (40 cycles); melting curve profile from 65 to 97 °C to verify the specificity of the reaction. The *A. citrulli GADPH* gene was used as an internal control to normalize gene expression. The threshold cycles (Ct) were determined with the Light Cycler 480 II software (Roche) and the fold-changes of three biological samples with three technical replicates per treatment were obtained by the ΔΔCt method [74]. Significant differences in expression values were evaluated using the Mann-Whitney non-parametrical test (α = 5%).

### Additional bioinformatics tools

BlastP analyses for search of T3E homologs were done at the NCBI server against the non-redundant protein sequences (nr) database, selecting the organisms *Acidovorax* (taxid: 12916), *Xanthomonas* (taxid: 338), *Ralstonia* (taxid: 48736) or *Pseudomonas syringae* group (taxid: 136849), with default parameters. Gene ontology (GO) assignments were done using Blast2GO software v5.2 (https://www.blast2go.com/). SignalP4.1 [75], Phobious [76] and Pred-Tat [77] were used for detection of N-terminal type II secretion signal peptides. The program fuzznuc (EMBOSS package; http://www.bioinformatics.nl/cgi-bin/emboss/fuzznuc) was used to detect perfect PIP box sequences (TTCGB-N15-TTCGB; [32]) in the *A. citrulli* M6 genome. A logo of the PIP box motif of *A. citrulli* M6 was done with MEME-ChiP [78] at the MEME Suite website (http://meme-suite.org/). Domain search of T3Es was carried out using the following databases/tools: Protein Data Bank (PDB) and UniProtKB/Swiss-Prot (through NCBI Blast), PFAM (https://pfam.xfam.org/), Prosite (https://prosite.expasy.org/) and InterPro (https://www.ebi.ac.uk/interpro/search/sequence-search). LogSidDB [79] and Plant-mPLoc [80] were used for detection of protein localization signals and for prediction of subcellular localization of T3Es, respectively.

### Translocation assays

The ORFs without the stop codon of candidate genes were amplified using specific primers (S7 Table) and cloned into the *SaT*I/*Xba*I sites of pBBR1MCS-*2::avrBs2_62-574_*, upstream to and in frame with the *avrBs2_62-574_* HR domain of *avrBs2* and an haemagglutinin (HA) tag [28], except for ORFs of genes *APS58_0500* and *APS58 1760*, which were cloned into the *Xho*I/*Xba*I sites of the same vector. The resulting plasmids were mobilized into *X. euvesicatoria* strains 85-10 *hrpG*AavrBs2* [81] and 85-10 *hrpG*ΔhrpF* [82]. Expression of recombinant T3E/CT3E-AvrBs2_62-574_-HA proteins was verified by Western blot using the iBlot Gel Transfer Stacks Nitrocellulose kit (Invitrogen), and anti-hemagglutinin (HA)-tag and horseradish peroxidase (HRP) antibodies (Cell Signaling Technology, Danvers, MA, USA) (S6 Fig). For translocation assays, *X. euvesicatoria* strains were grown overnight in LB broth with Km, centrifuged and resuspended in 10 mM MgCl2 to a concentration of 10^8^ cfu/mL. These suspensions were used to infiltrate the three youngest, fully expanded leaves of 5-week-old ECW20R and ECW30R [83] pepper plants, carrying and lacking the *Bs2* gene, respectively, using a needleless syringe. The plants were kept in a growth chamber at 25 °C, ~50% relative humidity, 12 h day/12 h night. HR was monitored 36 h after inoculation (h.a.i.). For visualization of cell death, the infiltrated leaves were treated as described above for pepper leaves infiltrated with *A. citrulli* strains. Each candidate gene was tested in three independent experiments with at least three plants, with similar results being obtained among replicates and experiments.

### *Agrobacterium-mediated* transient expression and confocal imaging

The ORFs of genes *APS58_0500, APS58_1448* and *APS58_4116* were amplified with specific primers (S7 Table) and cloned into pEarlyGate101 binary vector [84], upstream of a Yellow Fluorescence Protein (YFP) encoding gene and an HA tag using the Gateway cloning system (ThermoFisher Scientific). The resulting plasmids were verified by sequencing and mobilized into *A. tumefaciens* GV3101 as indicated above. Transient expression experiments were performed following the protocol described by Roden *et al*. [81] with few modifications. Briefly, overnight cultures of *A. tumefaciens*

GV3101 carrying the different plasmids were centrifuged, and pellets were resuspended in induction solution containing 10 mM MgCl2, 10 mM 2-(N-morpholino)-ethanesulfonic acid (MES), and 200 mM acetosyringone (pH 5.6). The suspensions were incubated at 25 °C without shaking for 3 h. Bacterial cultures were then diluted to OD_600nm_~0.6 and infiltrated with a needleless syringe into leaves of 4-week-old *N. benthamiana* plants [85] that were grown in a growth chamber (16 h/26 °C in the light, 8h/18°C in the dark; relative humidity set to 70%). Subcellular localization of tested T3Es coupled to YFP were investigated by co-infiltration with *A. tumefaciens* GV3101 carrying monomeric Red Fluorescence Protein fused in frame with the endoplasmid reticulum (ER) marker HDEL (mRFP-HDEL; [86, 87]), the membrane associated SlDRP2A (L. Pizarro and M. Bar, unpublished results) fused to monomeric Cherry fluorescent protein, and by staining with 1 mg/mL 4’,6-diamidino-2-phenylindole (DAPI), that was used to detect the nucleus of the plant cells [88]. As controls, plants were infiltrated with *A. tumefaciens* GV3101 carrying pEarlyGate104 (YFP-encoding gene). Infiltrated plants were kept in the growth chamber at similar conditions as above, and 48 h.a.i., functional fluorophores were visualized using a SPE (Leica Microsystems, Wetzlar, Germany) or a LSM 780 (Zeiss, Oberkochen, Germany) confocal microscope. Images were acquired using two tracks: track 1 for YFP detection, exciting at 514 nm and collecting emission from the emission range 530-560 nm; track 2 for RFP and mCherry detection, exciting at 561 nm and collecting from the emission range 588-641 nm. Images of 8 bits and 1024X1024 pixels were acquired using a pixel dwell time of 1.27, pixel averaging of 4 and pinhole of 1 airy unit. Analysis of colocalization was conducted with Fiji-ImageJ using the Coloc2 tool. For calculating the Pearson correlation coefficient, 15-18 images were analysed. Signal profiles were analysed using the Plot Profile tool [89].

## Acknowledgements

We thank Dr. Einat Zelinger from the Interdepartmental Core Facility of the Robert H. Smith Faculty of Agriculture, Food and Environment Hebrew University Interdepartmental Core Facility for her assistance with confocal microscopy. We also thank Dr. Inbar Plaschkes from the Bioinformatics Unit of the Hebrew University of Jerusalem for her assistance with the RNA-Seq data.

## Supporting information

**S1 Fig. HrpX and HrpG are required for pathogenicity of *Acidovorax citrulli* M6.**

Lesions induced in a melon (cv. HA61428) leaf syringe-infiltrated with 10^8^ cfu/mL suspensions of wild-type M6, but not of M6 mutants defective in *hrpX* and *hrpG* genes. Partial restoration of the wild-type phenotype was observed following transformation of the mutants with plasmids pBBR1MCS-5::hrpX and pBBR1MCS-5::hrpG (complementation plasmids), respectively. The picture was taken 3 days after infiltration.

**S2 Fig. Subcellular localization of APS_0500**. (A) Confocal microscopy images of *N. benthamiana* epidermal cells transiently expressing APS_0500-YFP and different endomembrane compartment markers as indicated. Representative images show APS_0500-YFP (green), the subcellular marker: HDEL-RFP, Free-mCherry or SlDRP2A (magenta) and the superimposed image of both channels (merge). Pearson correlation coefficient of the co-localization between APS_0500-YFP and the markers (N=15−18) was determined using the Coloc2 function from ImageJ. Data represented as mean ± SEM. (B) Confocal microscopy images of *N. benthamiana* epidermal cells transiently expressing the plasma membrane protein Flot1-GFP and Free-mCherry. All the images were acquired 48 h after *A. tumefaciens* infiltration using Zeiss LSM780 (40x/1,2 W Korr). Scale bar 20 μm.

**S3 Fig. Subcellular localization of APS_1448**. Confocal microscopy images of *N. benthamiana* epidermal cells transiently expressing APS_1448-YFP and different endomembrane compartment markers as indicated. Representative images show APS_1448-YFP (green), the subcellular markers HDEL-RFP, Free-mCherry or SlDRP2A (magenta), and the superimposed image of both channels (merge). Pearson correlation coefficient of the co-localization between APS_1448-YFP and the markers (N=15−18) was determined using the Coloc2 function from ImageJ. Data represented as mean ± SEM. All the images were acquired 48 h after *A. tumefaciens* infiltration using Zeiss LSM780 (40x/1,2 W Korr). Scale bar, 20 μm.

**S4 Fig. Subcellular localization of APS_4116**. Confocal microscopy images of *N. benthamiana* epidermal cells transiently expressing APS_1448-YFP and different endomembrane compartment markers as indicated. Representative images show APS_1448-YFP (green), the subcellular markers HDEL-RFP, Free-mCherry or SlDRP2A (magenta), and the superimposed image of both channels (merge). Pearson correlation coefficient of the co-localization between APS_1448-YFP and the markers (N=15−18) was determined using the Coloc2 function from ImageJ. Data represented as mean ± SEM. All the images were acquired 48 h after *A. tumefaciens* infiltration using Zeiss LSM780 (40x/1,2 W Korr). Scale bar, 20 μm.

**S5 Fig. Distribution of *Acidovorax citrulli* M6 type III effectors (T3Es) according to their amino acid length**. The data are from the annotation (GenBank accession CP029373) of the *A. citrulli* M6 ORFs.

**S6 Fig. Expression of effector-AvrBs262-574::HA fusion proteins of T3Es that were tested in translocation assays**. Total protein was extracted from overnight cultures of *Xanthomonas euvesicatoria 85-10-hrpG*-ΔavrBs2* expressing CT3E-AvrBs2_62-574_-HA fusions in plasmid *ęBBR1MCS-2::avrBs2_62_._514_*. Proteins were analysed by Western blot using HA-tag antibody. XopS *(X. euvesicatoria* effector)-AvrBs2_62-574_::HA was included as positive control. Asterisks indicate the size of the expected bands.

**S1 Table**. Ranking and prediction scores of open reading frames of *Acidovorax citrulli* AAC00-1 (GenBank accession CP000512.1) in the first machine learning run.

**S2 Table**. Occurrence of *Acidovorax citrulli* M6 type III effectors in other plant pathogenic *Acidovorax* species.

**S3 Table**. Differential gene expression as determined by RNA-Seq between *Acidovorax citrulli* M6 and an M6 mutant strain defective in *hrpX* gene, after 72 h of growth in XVM2 minimal medium at 28 °C.

**S4 Table**. Perfect plant-inducible promoter (PIP) boxes in the *Acidovorax citrulli* M6 genome.

**S5 Table**. Ranking and prediction scores of open reading frames of *Acidovorax citrulli* M6 (GenBank accession CP029373) in the second machine learning run.

**S6 Table**. Bacterial strains and plasmids used in this study.

**S7 Table**. DNA oligonucleotide primers used in this study.

**S8 Table**. List and description of the features used for the first and second machine learning runs.

