## Supplementary figures and images for "Show me your secret(ed) weapons: a multifaceted approach reveals novel type III-secreted effectors of a plant pathogenic bacterium"

### Supplemental Figure S1

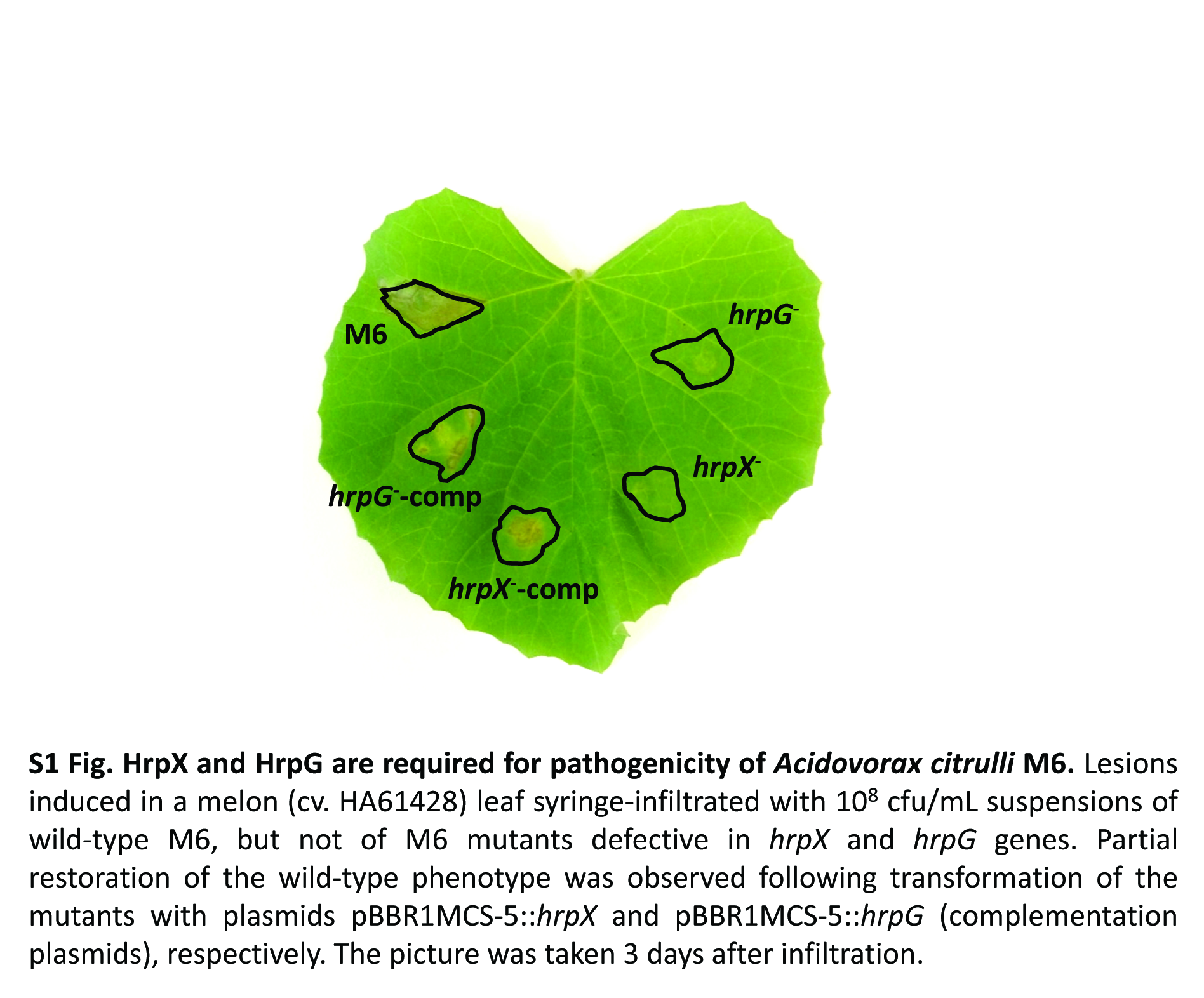

### Supplemental Figure S2

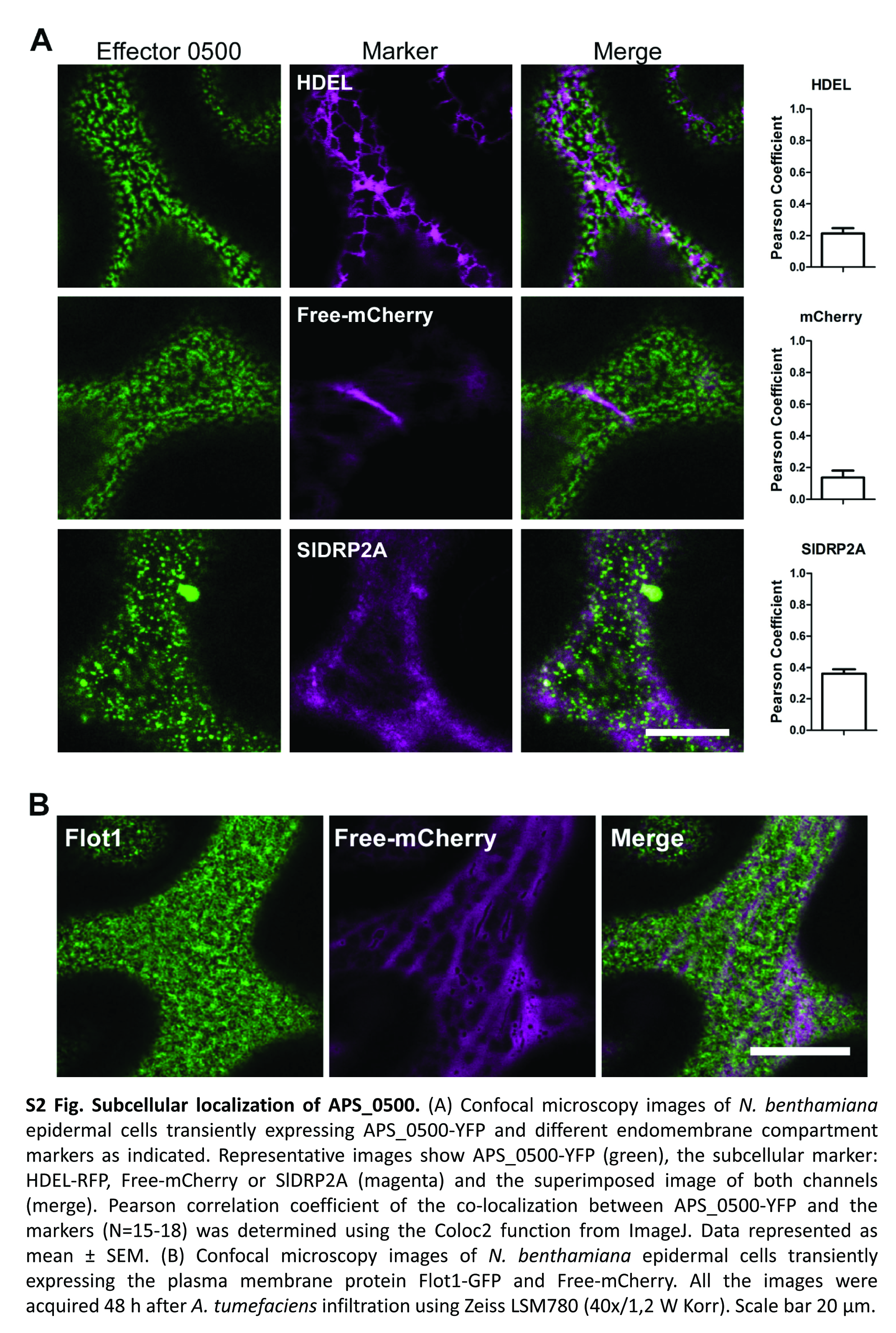

### Supplemental Figure S3

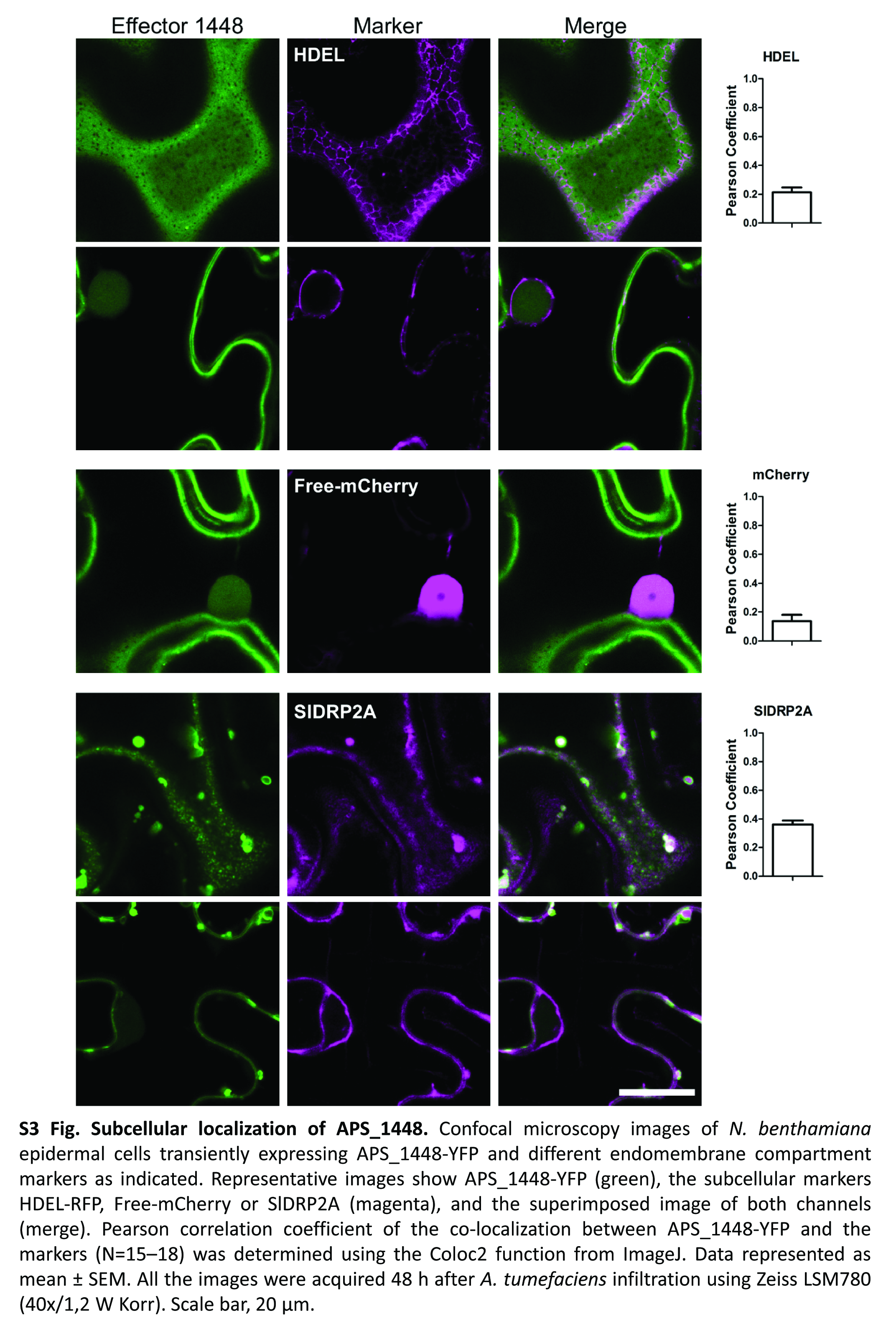

### Supplemental Figure S4

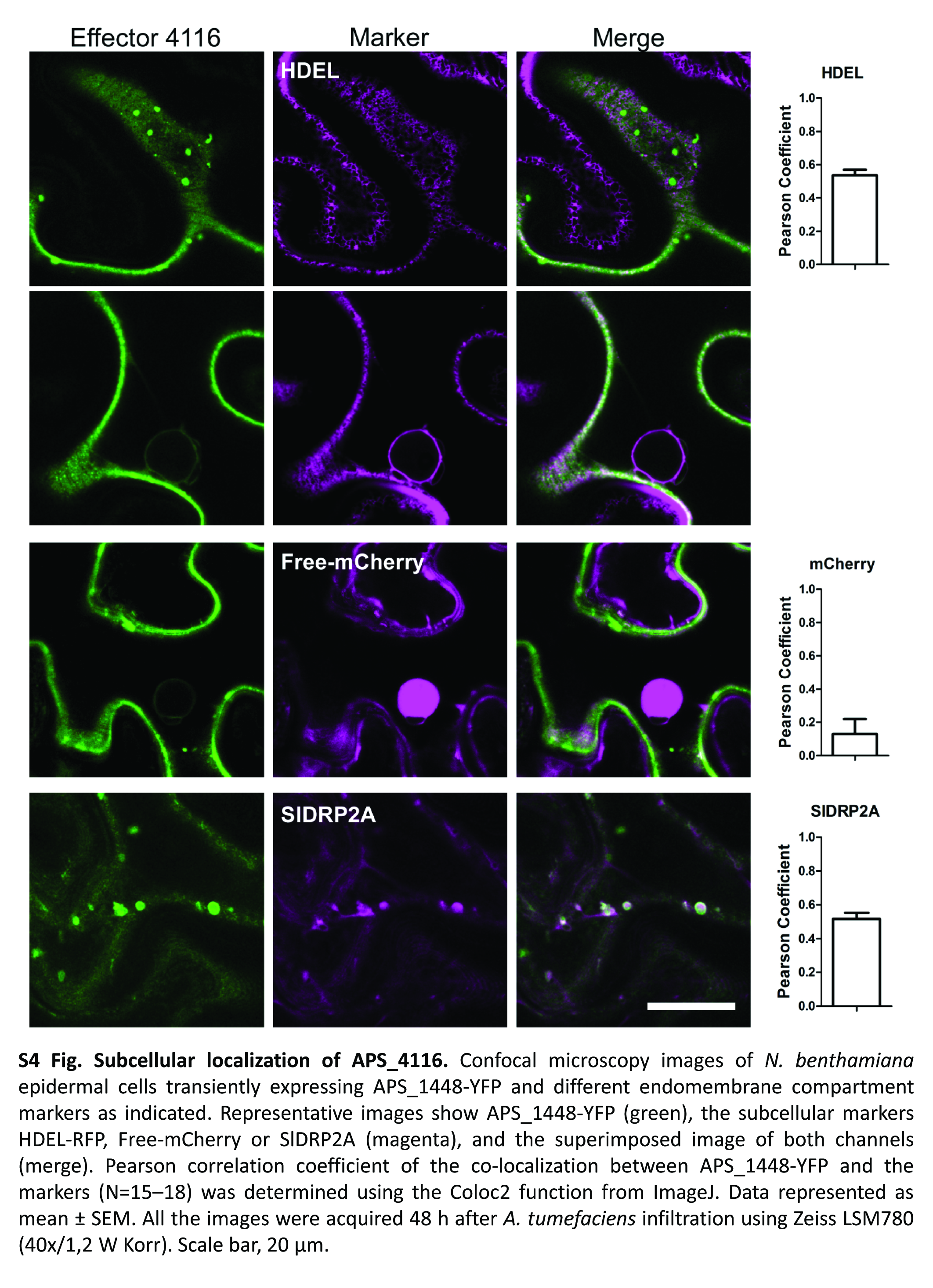

### Supplemental Figure S5

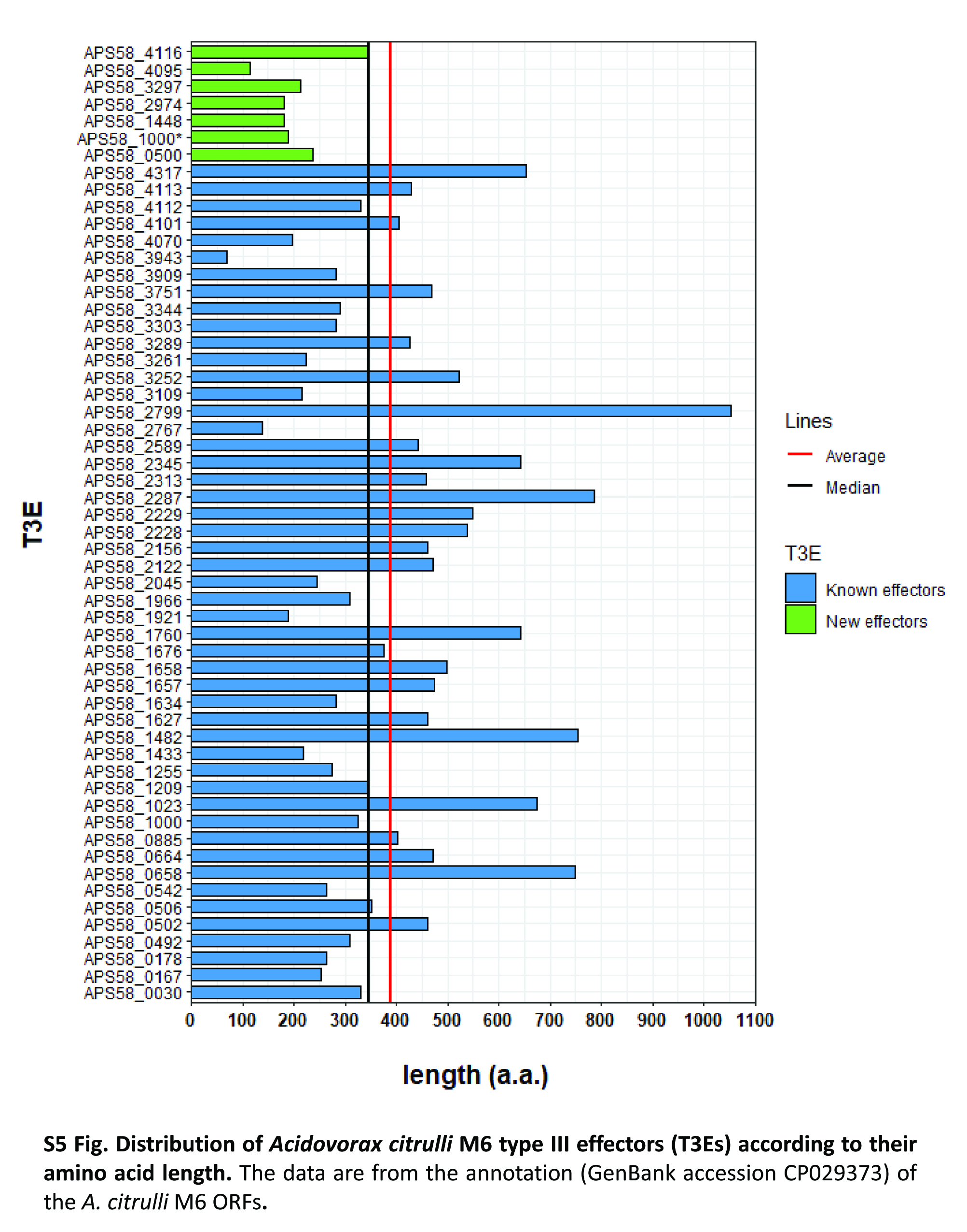

### Supplemental Figure S6

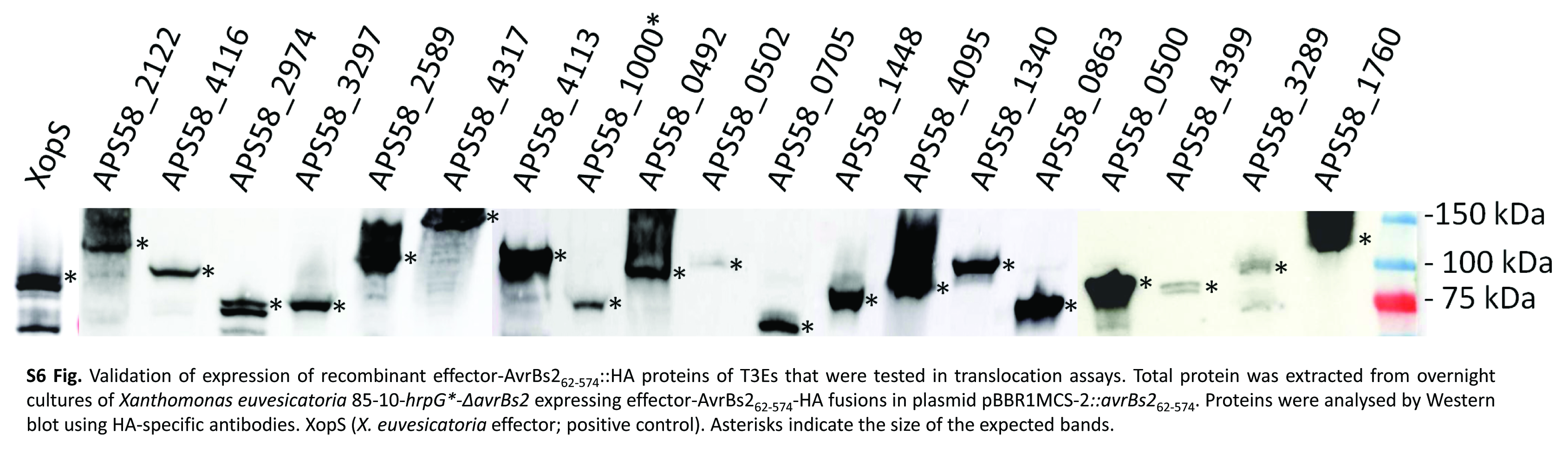
