## Supplemental Table S2 for "Show me your secret(ed) weapons: a multifaceted approach reveals novel type III-secreted effectors of a plant pathogenic bacterium"

**S2 Table. Occurrence of *Acidovorax citrulli* M6 type III effectors in other plant pathogenic *Acidovorax* species.**

| **T3Es similar to known effectors from other plant pathogenic bacteria (see Table 1)** | | | | | | | | |
| --- | --- | --- | --- | --- | --- | --- | --- | --- |
| **Gene ID in M6^1^** | **Locus tag in AAC00-1^2^** | ***Aci*^3^** | ***Aav*^3^** | ***Aor*^3^** | ***Aca*^3^** | ***Ako*^3^** | ***Aan*^3^** | ***Ava*^3^** |
| ***APS58_0030*** | *Aave_2531* | + | + | + | - | - | - | (+) |
| ***APS58_0167* ^4^** | *Aave_2166* | + | + | - | - | + | + | - |
| ***APS58_0178*** | *-* | + | + | + | + | - | - | + |
| ***APS58_0492*** | *Aave_3452* | + | + | + | + | + | + | + |
| ***APS58_0502*** | *Aave_3462* | + | + | + | + | - | (+) | - |
| ***APS58_0506*** | *-* | + | + | + | + | + | + | + |
| ***APS58_0542*** | *Aave_3502* | + | + | - | + | + | + | - |
| ***APS58_0658*** | *Aave_3621* | + | - | - | + | - | - | - |
| ***APS58_0664*** | *Aave_3626* | + | + | + | + | - | - | - |
| ***APS58_0885*** | *Aave_3847* | + | + | + | + | - | - | - |
| ***APS58_1000*** | *Aave_3961* | + | + | + | + | + | (+) | - |
| ***APS58_1023*** | *Aave_4359* | + | + | - | - | - | + | - |
| ***APS58_1209*** | *-* | + | + | (+) | (+) | (+) | + |  |
| ***APS58_1255*** | *Aave_4254* | + | + | + | (+) | + | - | - |
| ***APS58_1433*** | *Aave_4427* | + | - | - | - | - | - | - |
| ***APS58_1482*** | *Aave_4472* | + | + | + | + | + | + | (+) |
| ***APS58_1627*** | *Aave_4606* | + | + | + | + | + | + | - |
| ***APS58_1634*** | *Aave_4612* | + | + | + | + | + | + | - |
| ***APS58_1657*** | *Aave_4631* | + | + | + | + | + | + | + |
| ***APS58_1658*** | *Aave_4632* | + | + | + | + | + | + | + |
| ***APS58_1676*** | *-* | + | (+) | (+) | - | (+) | + | + |
| ***APS58_1760*** | *Aave_4728* | + | + | + | + | + | + | - |
| ***APS58_1921*** | *Aave_0085* | + | + | + | + | - | (+) | - |
| ***APS58_1966*** | *-* | + | + | + | - | + | + | - |
| ***APS58_2045*** | *Aave_0201* | + | + | + | - | - | - | - |
| ***APS58_2122*** | *Aave_0277* | + | + | - | - | - | - | - |
| ***APS58_2156*** | *Aave_0310* | + | + | + | + | - | - | - |
| ***APS58_2228*** | *-* | + | + | + | + | + | (+) | + |
| ***APS58_2229*** | *-* | + | + | + | + | - | - | - |
| ***APS58_2287*** | *Aave_0433* | + | + | - | + | - | - | - |
| ***APS58_2313*** | *Aave_0458* | + | + | + | + | + | + | + |
| ***APS58_2345*** | *Aave_0588* | + | + | + | (+) | + | (+) | + |
| ***APS58_2589*** | *Aave_0889* | + | + | + | (+) | - | - | - |
| ***APS58_2767*** | *-* | + | + | + | - | + | - | + |
| ***APS58_2799*** | *Aave_1090* | + | + | + | + | + | (+) | + |
| ***APS58_3109*** | *Aave_1373* | + | + | - | - | (+) | + | - |
| ***APS58_3252*** | *Aave_1508* | + | + | + | + | + | + | + |
| ***APS58_3261*** | *Aave_1520* | + | (+) | - | - | + | - | - |
| ***APS58_3289*** | *Aave_1548* | + | + | + | + | + | - | - |
| ***APS58_3303*** | *-* | + | + | + | + | + | + | + |
| ***APS58_3344*** | *Aave_1647* | + | + | - | - | - | - | - |
| ***APS58_3751*** | *Aave_3237* | + | + | + | + | + | + | + |
| ***APS58_3909*** | *Aave_3085* | + | + | + | + | - | - | - |
| ***APS58_3930*** | *Aave_3063* | + | + | - | - | - | - | - |
| ***APS58_3931* ^4^** | *Aave_3062* | + | + | - | - | - | - | - |
| ***APS58_3943*** | *-* | + | + | - | (+) | + | - | - |
| ***APS58_4070*** | *Aave_2876* | + | + | + | - | (+) | - | (+) |
| ***APS58_4101*** | *Aave_2844* | + | + | + | - | - | + | + |
| ***APS58_4112*** | *Aave_2173* | + | + | - | - | - | - | - |
| ***APS58_4113*** | *Aave_2174* | + | + | + | - | - | - | - |
| ***APS58_4317*** | *Aave_2802* | + | + | + | + | - | - | - |
| **T3Es discovered in this study with no homology to known effectors** | | | | | | | | |
| **Gene ID in M6^1^** | **Locus tag in AAC00-1^2^** | ***Aci*^3^** | ***Aav*^3^** | ***Aor*^3^** | ***Aca*^3^** | ***Ako*^3^** | ***Aan*^3^** | ***Ava*^3^** |
| ***APS58_0500*** | *Aave_3459* | + | + | + | + | - | - | - |
| ***APS58_1000**** ^5^ | *Aave_3960* | + | + | + | + | (+) | - | - |
| ***APS58_1448*** | *Aave_4442* | + | + | (+) | - | + | + | - |
| ***APS58_2974*** | *Aave_1244* | + | + | - | - | - | - | - |
| ***APS58_3297*** | *Aave_1555* | + | + | + | + | (+) | (+) | - |
| ***APS58_4095*** | *Aave_2850* | + | + | + | + | (+) | (+) | (+) |
| ***APS58_4116*** | *Aave_2177* | + | - | - | - | (+) | (+) | (+) |

^1^ Gene IDs are according to the recent annotation of the *A. citrulli* M6 chromosome (GenBank accession CP029373).

^2^ Corresponding locus_tag in the group II model strain of *A. citrulli*, AAC00-1 (GenBank accession CP000512.1). "-" means that this gene is not present in strain AAC00-1.

^3^ Detected (+) or non-detected (-) in other plant-pathogenic *Acidovorax* species as determined by BlastP against the non-redundant protein sequences (nr) database [*Acidovorax* taxid (12916)]: *Aci*, *A. citrulli* (7); *Aav*, *A. avenae* (18); *Aor*, *A. oryzae* (2); *Aca*, *A. cattleyae* (1); *Ako*, *A. konjaci* (1); *Aan*, *A. anthhurii* (1); and *Ava*, *A. valerianellae* (1). The numbers between parentheses indicate the number of genomes available in the database at the time this analysis was carried out.

(+) indicates significant similarity to hits with relatively low query coverage (below 60%).

^4^ These genes are probably non-functional in strain M6 and in all group I strains assessed so far (Eckshtain-Levi *et al*., 2014).

^5^ *APS58_1000**: this gene ranked high in the first ML but was not annotated in the recent annotation of *A. citrulli* M6. Its ORF is located between genes *APS58_0999* and *APS58_1000* (positions 1129817-1130383).
