## Supplemental Table S6 for "Show me your secret(ed) weapons: a multifaceted approach reveals novel type III-secreted effectors of a plant pathogenic bacterium"

**S6 Table. Bacterial strains and plasmids used in this study.**

| Strain/plasmid | Relevant properties | Source or reference |
| --- | --- | --- |
| ***Acidovorax citrulli*** | | |
| M6 | wild type, group I strain; Ap^R^ | Burdman *et al.*, 2005 |
| M6 *hrcV^-^* | M6 derivate disrupted in the *hrcV* gene; Ap^R^, Km^R^ | Bahar and Burdman, 2010 |
| M6 *hrpX^-^* | M6 derivate disrupted in the *hrpX* gene; Ap^R^, Km^R^ | This study |
| M6 *hrpG^-^* | M6 derivate disrupted in the *hrpG* gene; Ap^R^, Km^R^ | This study |
| M6 *hrpX^-^* (pBBR1MCS-5*::hrpX*) | M6 *hrpX*^-^ mutant complemented with pBBR1MCS-5*::hrpX*; Ap^R^, Km^R^ Gm^R^ | This study |
| M6 *hrpG^-^* (pBBR1MCS-5*::hrpG*) | M6 *hrpG*^-^ mutant complemented with pBBR1MCS-5*::hrpG*; Ap^R^, Km^R^ Gm^R^ | This study |
| ***Xanthomonas euvesicatoria*** | | |
| 85-10 *hrpG**∆*avrBs2* | 85-10 derivative containing the *hrpG** mutation (overexpressing this gene) and a deletion in *avrBs2*; Rif^R^, Gm^R^ | Roden *et al.*, 2004 |
| 85-10 *hrpG**∆*hrpF* | 85-10 derivative containing the *hrpG** mutation and a deletion in *hrpF*; Rif^R^ | Casper-Lindley *et al.*, 2002 |
| 85-10 *hrpG**∆*avrBs2* (pBBR1MCS-2::*avrBs2_62-574_*) | 85-10 *hrpG**∆*avrBs2* derivate carrying a plasmid with the AvrBs2 HR domain without the N-terminal translocation signal (AvrBs2_62-574_); negative control in translocation assays; Rif^R^, Km^R^ | Teper *et al.*, 2015 |
| 85-10 *hrpG**∆*avrBs2* (pBBR1MCS-2*::xopS::avrBs2_62-574_*) | 85-10 *hrpG**∆*avrBs2* derivate carrying a plasmid with the *X. euvesicatoria* 85-10 *xopS* ORF without the stop codon fused in frame with AvrBs2_62-574_; positive control in translocation assays; Rif^R^, Km^R^ | Teper *et al.*, 2015 |
| 85-10 *hrpG**∆*avrBs2* (pBBR1MCS-2*::APS58_XXXX::avrBs2_62-574_*) | Nineteen 85-10 *hrpG**∆*avrBs2* derivates carrying a plasmid with an *APS58_XXXX* ORF without the start codon fused in frame with AvrBs2_62-574_; *XXXX* refers to the gene ID of *A. citrulli* M6 genes tested in translocation assays; Rif^R^, Km^R^ | This study |
| 85-10 *hrpG**∆*hrpF* (pBBR1MCS-2*::APS58_XXXX::avrBs2_62-574_*) | Nineteen 85-10 *hrpG**∆*hrpF* derivates carrying a plasmid with an *APS58_XXXX* ORFs without the start codon fused in frame with AvrBs2_62-574_; *XXXX* refers to the gene ID of *A. citrulli* M6 genes tested in translocation assays; Rif^R^, Km^R^ | This study |
| ***Agrobacterium tumefaciens*** | | |
| GV3101 | wild type; Rif^R^ | Rotino and Gleddie, 1990 |
| ***Escherichia coli*** | | |
| DB3.1 | *gyrA462*, *endA1*, ∆(*sr1-recA*), *mcrB*, *mrr*, *hsdS20*, *glnV44* (=*supE44*), *ara14*, *galK2*, *lacY1*, *proA2*, *rpsL20*, *xyl5*, *leuB6*, *mtl1* | Invitrogen (Carlsbad, California) |
| DH5α | *supE44*, ∆*lacU169*, *hsdR17*, *recA1*, *endA1*, *gyrA96*, *thi-1*, *relA1*, Nx^R^ | Sambrook *et al.*, 1989 |
| S17-1 λpir | ∆lysogenic S17-1 derivate producing π protein for replication of plasmids carrying *oriR6K*; *recA*, *pro*, *hsdR,* RP4-2-Tc::Mu-Km::Tn7, λ-pir | Simon *et al.*, 1983 |
| **Plasmids** | | |
| pJP5603 | R6K-based suicide vector; requires the pir-encoded π protein for replication; used for mutagenesis of *A. citrulli*; Km^R^ | Penfold and Pemberton, 1992 |
| pBBR1MCS-5 | P_T7_ rep broad host expression vector; Gm^R^ | Kovach *et al.*, 1995 |
| pJP5603::*hrpX*_int_ | pJP5603 carrying an internal fragment (383 bp) of *hrpX* with an early stop codon, inserted into the *Bam*HI/*Eco*RI sites; used for generation of *A. citrulli* M6 *hrpX*^-^ mutant by insertional mutagenesis following single homologous recombination; Km^R^ | This study |
| pBBR1MCS-5::*hrpX* | pBBR1MCS-5 carrying the *hrpX* ORF (1407 bp) inserted into the *Eco*RI/*Bam*HI sites and under the control of the lacZα promoter; used for complementation of the *A. citrulli* M6 *hrpX*^-^ mutant; Gm^R^ | This study |
| pJP5603*::hrpG*_int_ | pJP5603 carrying an internal fragment (438 bp) of *hrpG* with an early stop codon, inserted into the *Bam*HI/*Eco*RI sites; used for generation of *A. citrulli* M6 *hrpG*^-^ mutant by insertional mutagenesis following single homologous recombination; Km^R^ | This study |
| pBBR1MCS-5*::hrpG* | pBBR1MCS-5 carrying the *hrpG* ORF (801 bp) inserted into the *Eco*RI/*Bam*HI sites and under the control of the lacZα promoter; used for complementation of the *A. citrulli* M6 *hrpG*^-^ mutant; Gm^R^ | This study |
| pDONR207 | Gateway donor vector; Gm^R^ | Invitrogen |
| pEarleyGate100 | Gateway-compatible plant transformation vector; Km^R^ | Earley *et al.*, 2006 |
| pEarleyGate101 | Gateway-compatible plant transformation vector with YFP and HA C-terminal tags; Km^R^ | Earley *et al.,* 2006 |
| pEarleyGate104 | Gateway-compatible plant transformation vector with a YFP N-terminal tag; Km^R^ | Earley *et al.*, 2006 |
| pDONR207*::APS58_0500* | pDONR207 carrying the *APS58_0500* ORF without the stop codon (711 bp); Gm^R^ | This study |
| pEarleyGate101*::APS58_0500* | pEarleyGate101 carrying the *APS58_0500* ORF without the stop codon fused to YFP and HA; Km^R^ | This study |
| pDONR207*::APS58_1448* | pDONR207 carrying the *APS58_1448* ORF without the stop codon (543 bp); Gm^R^ | This study |
| pEarleyGate101*::APS58_1448* | pEarleyGate101 carrying the *APS58_1448* ORF without the stop codon fused to YFP and HA; Km^R^ | This study |
| pDONR207*::APS58_4116* | pDONR207 carrying the *APS58_4116* ORF without the stop codon (1029 bp); Gm^R^ | This study |
| pEarleyGate101*::APS58_4116* | pEarleyGate101 carrying the *APS58_4116* ORF without the stop codon fused to YFP and HA; Km^R^ | This study |
| pmRFP-HDEL | Plasmid used for transient expression of the endoplasmic reticulum marker HDEL fused to red fluorescent protein (RFP); Km^R^ | Runions *et al*., 2006; Schoberer *et al*., 2009 |
| pBIN20::ER‐rk (CD3‐959) | pBIN20 carrying HDEL endoplasmic reticulum retention peptide fused to mCherry; Km^R^ | Nelson *et al.*, 2007 |
| pCAMBIA1200::GFP-Flot1 | pCAMBIA1200 carrying the plasma membrane protein Flot1 from *Arabidopsis thaliana* fused to green fluorescent protein (GFP) under the CaMV 35S promoter; Km^R^ | Li *et al.*, 2012 |
| pBINPLUS::SlDRP2A-mCherry | pBINPLUS carrying the Dynamin SlDRP2A from *Solanum lycopersicum* fused to mCherry under the CaMV 35S promoter; Km^R^ | L. Pizarro and M. Bar, unpublished |
| pBINPLUS::mCherry | pBINPLUS carrying mCherry under the CaMV 35S promoter; Km^R^ | Leibman-Markus *et al.*, 2018. |
| pBBR1MCS-2*::avrBs2_62-574_* | pBBR1MCS-2 carrying the region encoding the HR-inducing domain of AvrBs2, AvrBs2_62-574_, inserted into the *Xba*I/*Sac*I sites and upstream of an HA tag; Km^R^ | Teper *et al.*, 2015 |
| pBBR1MCS-2*::APS58_XXXX::avrBs2_62-574_* | pBBR1MCS-2*::avrBs2_62-574_* carrying *APS58_XXXX* ORFs without the stop codon, inserted into the *Sal*I/*Xba*I sites upstream and in frame with *avrBs2_62-574_*; used to assess translocation; seventeen constructs with *XXXX* being *0492*, *0502*, *0705*, *0863*, *1000** (non-annotated gene; located between genes *APS58_0999* and *APS58_1000*), *1340*, *1448*, *2122*, *2589*, *2974*, *3289*, *3297*, *4095*, *4113*, *4116*, *4317*, *4399*; Km^R^ | This study |
| pBBR1MCS-2*::APS58_XXXX::avrBs2_62-574_* | pBBR1MCS-2*::avrBs2_62-574_* carrying *APS58_XXXX* ORFs without the stop codon, inserted into the *Xho*I/*Xba*I sites upstream and in frame with *avrBs2_62-574_*; used to assess translocation; two constructs with *XXXX* being *0500* and *1760*; Km^R^ | This study |

^*^ Ap^R^, Gm^R^, Km^R^ and Rif^R^ indicate resistance to ampicillin, gentamicin, kanamycin and rifampicin, respectively.
