## Supplemental Table S7 for "Show me your secret(ed) weapons: a multifaceted approach reveals novel type III-secreted effectors of a plant pathogenic bacterium"

**S7 Table. DNA oligonucleotide primers used in this study.**

| Name | Sequence^1^ | Usage |
| --- | --- | --- |
| pJP_F | 5'-CAGGAAACAGCTATGACCAT-3' | Sequence verification of inserts cloned into pJP5503 |
| pJP_R | 5'-GATTAAGTTGGGTAACGCCA-3' |  |
| pBBR_F | 5'-TGTGGAATTGTGAGCGGATA-3' | Sequence verification of inserts cloned into pBBR1MCS plasmids |
| pBBR_R | 5'-GTTTTCCCAGTCACGACGTT-3' |  |
| hrpXM6_mut_BamHI_F | 5'-AAAGGATCCAAGAACTGT**A**GGACCACG-3' | Used for amplification of an internal fragment of *APS58_2298* (*hrpX*) for mutagenesis |
| hrpXM6_mut_EcorI_R | 5'-AAAGAATTCCGACAG**T**CAGTACTGGAT-3' |  |
| hrpXM6_comp_EcoRI_F | 5'-AAAGAATTCATGCTCCTCTCTCTGCTTTCTC-3' | Used for amplification of the complete ORF of *APS58_2298* (*hrpX*), for complementation and for verification of the *hrpX* mutant |
| hrpXM6_comp_BamHI_R | 5'-ATAGGATCCTCAGTGCCGCATCGACGACAGC -3' |  |
| hrpGM6_mut_BamHI_F | 5'-ATAGGATCCGGATCTGGGCCTGATC**T**GAA-3' | Used for amplification of an internal fragment of *APS58_2299* (*hrpG*) for mutagenesis |
| hrpGM6_mut_EcoRI_R | 5'-AAAGAATTCCATCCGCACTGGCGCAGCT**A**-3' |  |
| hrpGM6_comp_EcoRI_F | 5'-ATAGAATTCATGCGAGTCGCATTGCTGT-3' | Used for amplification of the complete ORF of *APS58_2299* (*hrpG*), for complementation and for verification of the *hrpG* mutant |
| hrpGM6_comp_BamHI_R | 5'-ATAGGATCCTCAACTGCCCAGGGGCGGCA-3' |  |
| APS58_2306-RT_F | 5'-TCATCACGCTGGTCAACATC-3' | Used to assess expression of *APS58_2306* (*hrcV*) gene by RT-PCR |
| APS58_2306-RT_R | 5'- GGCCATACAGATGAAGAGCG-3' |  |
| APS58_2309-RT_F | 5'- CTGATCGGATTCGCCGTC-3' | Used to assess expression of *APS58_2309* (*hrcT*) gene by RT-PCR and by qRT-PCR |
| APS58_2309-RT_R | 5'- CCAATTGCATGAGCAGGTTG-3' |  |
| APS58_2321-RT_F | 5'- GAACGTGCAGGTGGAAAAGA-3' | Used to assess expression of *APS58_2321* (*hrcJ*) gene by RT-PCR |
| APS58_2321-RT_R | 5'- GGCGACGCCATAGATGAAG-3' |  |
| APS58_2331-RT-F | 5'- TATTTCTACCCGGGCAAGTC-3' | Used to assess expression of *APS58_2331* (*hrcC*) gene by RT-PCR |
| APS58_2331-RT-R | 5'- GCCCGGAAACATAAATCGTG-3' |  |
| APS58_3289-RT_F | 5'- GGATTTCATCCGCTTCATGG-3' | Used to assess expression of *APS58_3289* (*hopW1-1*) gene by RT-PCR and by qRT-PCR |
| APS58_3289-RT_R | 5'- TCGCACATCAATGACGGAG-3' |  |
| APS58_1610-RT_F | 5'- TCGGCCGTGCAGAACTTC-3' | Used to assess expression of *APS58_1610 (GAPDH)* gene by RT-PCR |
| APS58_1610-RT_R | 5'- TGCAGCATGTAGGCCAGGTA-3' |  |
| APS58_0502q_F | 5'-GTGGTGCACATGAACTCGT-3' | Used to assess expression of *APS58_0502* (*yopJ*) gene by qRT-PCR |
| APS58_0502q_R | 5'-CGACAGCGCGAAGATCC-3' |  |
| APS58_1340q_F | 5'-GAGACCGCCAAGGAATACG-3' | Used to assess expression of *APS58_1340* gene by qRT-PCR |
| APS58_1340q_R | 5'-CCGATCGAGGTGTTGTTGTC-3' |  |
| APS58_2316q_F | 5'-CTGATGGTGAACGTGGGTTT-3' | Used to assess expression of *APS58_2316* (*hpaH*) gene by qRT-PCR |
| APS58_2316q_R | 5'-GATCATGCCGAACTTGGACT-3' |  |
| APS58_2331q_F | 5'-TATTTCTACCCGGGCAAGTC-3' | Used to assess expression of *APS58_2331* (*hrcC*) gene by qRT-PCR |
| APS58_2331q_R | 5'-GCCCGGAAACATAAATCGTG-3' |  |
| APS58_2334q_F | 5'-CTTTTCCCGCGGCATGAAT-3' | Used to assess expression of *APS58_2334* gene by qRT-PCR |
| APS58_2334q_R | 5'-CGGTACTGGCGAACACG-3' |  |
| APS58_2764q_F | 5'- GATGTCGGCCTTCCTGATCG-3' | Used to assess expression of *APS58_2734* (*tatC*) gene by qRT-PCR |
| APS58_2764q_R | 5'- AGAAGTAGCAGAACGCCACG-3' |  |
| APS58_2974q_F | 5'-CAACAGGAGCTTCACAGTCT-3' | Used to assess expression of *APS58_2974* gene by qRT-PCR |
| APS58_2974q_R | 5'-TTGATAGACCACCTCGGGTT-3' |  |
| APS58_4116q_F | 5'-CATGACTACCCATGCGGC-3' | Used to assess expression of *APS58_4116* gene by qRT-PCR |
| APS58_4116q_R | 5'-CATCGTCAATGGCGTCGATA-3' |  |
| APS58_4317q_F | 5'- GACTACGTACAGGCGAAGC-3' | Used to assess expression of *APS58_4317* gene by qRT-PCR |
| APS58_4317q_R | 5'- CCAGGTGAATTTCCCGTACC-3' |  |
| APS58_0492_SalI_F | 5'-AAAGTCGACGTGGCGGCCGGTGCCGAC-3' | Used for cloning of the *APS58_0492* ORF without the stop codon in plasmid pBBR1MCS-2*::avrBs2_62-574_* |
| APS58_0492_XbaI_R | 5'-AAATCTAGAGGACGTTCTCCGGCGGAG-3' |  |
| APS58_0500_XhoIF | 5'-AAACTCGAGATGCATTTCAGCTTCAATCA-3' | Used for cloning of the *APS58_0500* ORF without the stop codon in plasmid pBBR1MCS-2*::avrBs2_62-574_* |
| APS58_0500_XbaIR | 5'-AAATCTAGAGAAAACGTACTTGCCCAGC-3' |  |
| APS58_0502_SalI_F | 5'-AAAGTCGACATGCCGTCGCCGCCGCGG-3' | Used for cloning of the *APS58_0502* ORF without the stop codon in plasmid pBBR1MCS-2*::avrBs2_62-574_* |
| APS58_0502_XbaI_R | 5'-AAATCTAGAGTGCCGATACCAGTCGCG-3' |  |
| APS58_0705_SalI_F | 5'-AAAGTCGACCCATGAGCGACAACACCCAGC-3' | Used for cloning of the *APS58_0705* ORF without the stop codon in plasmid pBBR1MCS-2*::avrBs2_62-574_* |
| APS58_0705_XbaI_R | 5'-AAATCTAGAGGCCTGTGTCTGGCCGC-3' |  |
| APS58_0863_SalI_F | 5'-AAAGTCGACCGCCCCGATCCGCCTTTCCT-3' | Used for cloning of the *APS58_0863* ORF without the stop codon in plasmid pBBR1MCS-2*::avrBs2_62-574_* |
| APS58_0863_XbaI_R | 5'-AAATCTAGAGGCGCCGCCGATGCCGA-3' |  |
| APS58_1000*_SalI_F | 5'- AAAGTCGACAACATGCGCTGCGCCCCC -3' | Used for cloning of the *APS58_1000** ORF without the stop codon in plasmid pBBR1MCS-2*::avrBs2_62-574_* |
| APS58_1000*_XbaI_R | 5'- AAATCTAGACGCTGGCATGGGAATAGTCCC -3' |  |
| APS58_1340_SalI_F | 5'-AAAGTCGACCACCGGGCGCATCGACGC-3' | Used for cloning of the *APS58_1340* ORF without the stop codon in plasmid pBBR1MCS-2*::avrBs2_62-574_* |
| APS58_1340_XbaI_R | 5'-AAATCTAGAGCGGCCCGGGTTCATCAC-3' |  |
| APS58_1448_SalI_F | 5'-AAAGTCGACATGAAGCCCGTAGGAAATTT-3' | Used for cloning of the *APS58_1448* ORF without the stop codon in plasmid pBBR1MCS-2*::avrBs2_62-574_* |
| APS58_1448_XbaI_R | 5'-AAATCTAGAGCGGTCACTCAGCTGGA-3' |  |
| APS58_1760_XhoI_F | 5'-AAACTCGAGATGTCTGCGATCAACAGTTC-3' | Used for cloning of the *APS58_1760* ORF without the stop codon in plasmid pBBR1MCS-2*::avrBs2_62-574_* |
| APS58_1760_XbaI_R | 5'-AAATCTAGAGGGGGCCACGGGCGCGTG-3' |  |
| APS58_2122_SalI_F | 5'-AAAGTCGACATGAGAAAAGAACTAGCCAG-3' | Used for cloning of the *APS58_2122* ORF without the stop codon in plasmid pBBR1MCS-2*::avrBs2_62-574_* |
| APS58_2122_XbaI_R | 5'-AAATCTAGAGGCAGGCCTGCCCGGCAG-3' |  |
| APS58_2589_SalI_F | 5'-AAAGTCGACTTGTACCGCATAGGCAGG-3' | Used for cloning of the *APS58_2589* ORF without the stop codon in plasmid pBBR1MCS-2*::avrBs2_62-574_* |
| APS58_2589_XbaI_R | 5'-AAATCTAGACAGCGTGTAGGGGTCTGG-3' |  |
| APS58_2974_SalI_F | 5'-AAAGTCGACATGGGAAATCAATGCATCGG-3' | Used for cloning of the *APS58_2974* ORF without the stop codon in plasmid pBBR1MCS-2*::avrBs2_62-574_* |
| APS58_2974_XbaI_R | 5'-AAATCTAGAACGCCAGTCGCTGAACTG-3' |  |
| APS58_3289_SalI_F | 5'-AAAGTCGACATGCCTCTACAGTTCATTTC-3' | Used for cloning of the *APS58_3289* ORF without the stop codon in plasmid pBBR1MCS-2*::avrBs2_62-574_* |
| APS58_3289_XbaI_R | 5'-AAATCTAGATGGTTGATCCCCCGTCCGA-3' |  |
| APS58_3297_SalI_F | 5'-AAAGTCGACATGACCGATGTCCGGTTC-3' | Used for cloning of the *APS58_3297* ORF without the stop codon in plasmid pBBR1MCS-2*::avrBs2_62-574_* |
| APS58_3297_XbaI_R | 5'-AAATCTAGAGAAGAAGTGGGGGAAGCTCA-3' |  |
| APS58_4095_SalI_F | 5'-AAAGTCGACATGAACGCACGCACCGGC-3' | Used for cloning of the *APS58_4095* ORF without the stop codon in plasmid pBBR1MCS-2*::avrBs2_62-574_* |
| APS58_4095_XbaI_R | 5'-AAATCTAGAGGTGACCAGATCGGCCAC-3' |  |
| APS58_4113_SalI_F | 5'-AAAGTCGACATGTTGCCTGAATCCATC-3' | Used for cloning of the *APS58_4113* ORF without the stop codon in plasmid pBBR1MCS-2*::avrBs2_62-574_* |
| APS58_4113_XbaI_R | 5'-AAATCTAGAGGAGACGGGGCTCAGCAG-3' |  |
| APS58_4116_SalI_F | 5'-AAAGTCGACATGAAAAACTCCATCGGC-3' | Used for cloning of the *APS58_4116* ORF without the stop codon in plasmid pBBR1MCS-2*::avrBs2_62-574_* |
| APS58_4116_XbaI_R | 5'-AAATCTAGAGGGGCGGCGGCGGGCTTC-3' |  |
| APS58_4317_SalI_F | 5'-AAAGTCGACATGACCTTCCTGCAATTTCCCGC-3' | Used for cloning of the *APS58_4317* ORF without the stop codon in plasmid pBBR1MCS-2*::avrBs2_62-574_* |
| APS58_4317_XbaI_R | 5'-AAATCTAGACACCGGCGCGGAATGGCG-3' |  |
| APS58_4399_SalI_F | 5'-AAAGTCGACATGCCCGATACCACCTCTTCCG-3' | Used for cloning of the *APS58_4399* ORF without the stop codon in plasmid pBBR1MCS-2*::avrBs2_62-574_* |
| APS58_4399_XbaI_R | 5'-AAATCTAGAGGCCTTGCCGTCGGCGC-3' |  |
| 0500M6_attB1 | 5'-GGGGACAAGTTTGTACAAAAAAGCAGGCTTAATGCATTTCAGCTTCAAT-3' | Used for cloning of the *APS58_0500* ORF without the stop codon in plasmid pDONR207 by the Gateway system |
| 0500M6ns_attB2 | 5'-GGGGACCACTTTGTACAAGAAAGCTGGGTAGAAAACGTACTTGCCCAG-3' |  |
| 1448M6_attB1 | 5'-GGGGACAAGTTTGTACAAAAAAGCAGGCTTAATGAAGCCCGTAGGAAAT-3' | Used for cloning of the *APS58_1448* ORF without the stop codon in plasmid pDONR207 by the Gateway system |
| 1448M6ns_attB2 | 5'-GGGGACCACTTTGTACAAGAAAGCTGGGTAGCGGTCACTCAGCTGGAC-3' |  |
| 4116M6_attB1 | 5'- GGGGACAAGTTTGTACAAAAAAGCAGGCTTAATGAAAAACTCCATCGGC -3' | Used for cloning of the *APS58_4116* ORF without the stop codon in plasmid pDONR207 by the Gateway system |
| 4116M6ns_attB2 | 5'- GGGGACCACTTTGTACAAGAAAGCTGGGTAGGGGCGGCGGCGGGCTTC -3' |  |

^1^ Underlined nucleotides indicate the restriction sites of the corresponding enzymes or the attB1 or attB2 sequences as indicated in the primer names. Bolded nucleotides in primers used for mutagenesis of *hrpX* and *hrpG* indicate substitutions relative to the original sequence for addition of an early stop codon.
