## Supplemental Table S8 for "Show me your secret(ed) weapons: a multifaceted approach reveals novel type III-secreted effectors of a plant pathogenic bacterium"

**S8 Table. List and description of the features used for the first and second machine learning runs.**

| **Feature description** | **Feature name** | **index** |
| --- | --- | --- |
| Computes the GC content in the ORF. | GC_content | **1** |
| Computes the length of the gene product. | protein_length | **2** |
| Given a vector of the average amino acids profile of the known effector, and a similar vector for the non-effectors, it computes the difference between the Euclidean distance from the average aa profile of the known effectors, and the Euclidean distance from the average aa profile of the non-effectors. | similarity_to_the_average_aa_profile_of_effectors_vs_non_effectors | **3** |
| Computes the portion of A in the entire protein. | A_in_pep | **4** |
| Computes the portion of A in the first 25 aa of the protein. | A_in_the_N-terminal_region | **5** |
| Computes the portion of C in the entire protein. | C_in_pep | **6** |
| Computes the portion of C in the first 25 aa of the protein. | C_in_the_N-terminal_region | **7** |
| Computes the portion of D in the entire protein. | D_in_pep | **8** |
| Computes the portion of D in the first 25 aa of the protein. | D_in_the_N-terminal_region | **9** |
| Computes the portion of E in the entire protein. | E_in_pep | **10** |
| Computes the portion of E in the first 25 aa of the protein. | E_in_the_N-terminal_region | **11** |
| Computes the portion of F in the entire protein. | F_in_pep | **12** |
| Computes the portion of F in the first 25 aa of the protein. | F_in_the_N-terminal_region | **13** |
| Computes the portion of G in the entire protein. | G_in_pep | **14** |
| Computes the portion of G in the first 25 aa of the protein. | G_in_the_N-terminal_region | **15** |
| Computes the portion of H in the entire protein. | H_in_pep | **16** |
| Computes the portion of H in the first 25 aa of the protein. | H_in_the_N-terminal_region | **17** |
| Computes the portion of I in the entire protein. | I_in_pep | **18** |
| Computes the portion of I in the first 25 aa of the protein. | I_in_the_N-terminal_region | **19** |
| Computes the portion of K in the entire protein. | K_in_pep | **20** |
| Computes the portion of K in the first 25 aa of the protein. | K_in_the_N-terminal_region | **21** |
| Computes the portion of L in the entire protein. | L_in_pep | **22** |
| Computes the portion of L in the first 25 aa of the protein. | L_in_the_N-terminal_region | **23** |
| Computes the portion of M in the entire protein. | M_in_pep | **24** |
| Computes the portion of M in the first 25 aa of the protein. | M_in_the_N-terminal_region | **25** |
| Computes the portion of N in the entire protein. | N_in_pep | **26** |
| Computes the portion of N in the first 25 aa of the protein. | N_in_the_N-terminal_region | **27** |
| Computes the portion of P in the entire protein. | P_in_pep | **28** |
| Computes the portion of P in the first 25 aa of the protein. | P_in_the_N-terminal_region | **29** |
| Computes the portion of Q in the entire protein. | Q_in_pep | **30** |
| Computes the portion of Q in the first 25 aa of the protein. | Q_in_the_N-terminal_region | **31** |
| Computes the portion of R in the entire protein. | R_in_pep | **32** |
| Computes the portion of R in the first 25 aa of the protein. | R_in_the_N-terminal_region | **33** |
| Computes the portion of S in the entire protein. | S_in_pep | **34** |
| Computes the portion of S in the first 25 aa of the protein. | S_in_the_N-terminal_region | **35** |
| Computes the portion of T in the entire protein. | T_in_pep | **36** |
| Computes the portion of T in the first 25 aa of the protein. | T_in_the_N-terminal_region | **37** |
| Computes the portion of V in the entire protein. | V_in_pep | **38** |
| Computes the portion of V in the first 25 aa of the protein. | V_in_the_N-terminal_region | **39** |
| Computes the portion of W in the entire protein. | W_in_pep | **40** |
| Computes the portion of W in the first 25 aa of the protein. | W_in_the_N-terminal_region | **41** |
| Computes the portion of Y in the entire protein. | Y_in_pep | **42** |
| Computes the portion of Y in the first 25 aa of the protein. | Y_in_the_N-terminal_region | **43** |
| Computes the average hydrophilicity of the first 25 aa of the protein, using aa index BLAS910101. | hydrophilicity_of_N-terminal_region_BLAS910101 | **44** |
| Computes the average amphiphilicity of the first 25 aa of the protein, using aa index MITS020101. | amphiphilicity_of_N-terminal_region_MITS020101 | **45** |
| Computes the average hydrophobicity of the first 25 aa of the protein, using aa index KUHL950101. | hydrophobicity_of_N-terminal_region_KUHL950101 | **46** |
| Computes the average hydrophobicity of the first 25 aa of the protein, using aa index CIDH920105. | hydrophobicity_of_N-terminal_region_CIDH920105 | **47** |
| A binary feature that checks for the existence of a complete PIP-box in the promoter of the gene.  In the 1^st^ ML: TTCGC-N15-TTCGC.  In the 2^nd^ ML: TTCGB-N15-TTCGB, B being any nucleotide except adenine_._ | existence_of_pip_box_full_upstream_to_AUG | **48** |
| A binary feature that checks for the existence of a PIP-box with up to one mismatch in the promoter of the gene.  The patterns used are like in feature number 48. | existence_of_pip_box_one_mismatch_upstream_to_AUG | **49** |
| A binary feature that checks for the existence of a complete TATA box at -300.  TATAAT. | existence_of_TATA_box_300bp_upstream_to_AUG | **50** |
| a binary feature that checks for the existence of SesYEG secretion signal in the protein, using SignalP version 4.1. | SecYEG_secretion_existence_(SignalP) | **51** |
| Checks for the D score of SignalP version 4.1, of the protein. | SecYEG_secretion_score_(SignalP) | **52** |
| Checks for homology to phytopathogens, and returns the bit score of the best homolog (cut off E.val = 0.01). | homology_to_phytopatogens_(best_bit_score) | **53** |
| Checks for homology to phytopathogens, and returns the number of homologs (cut off E.val = 0.01). | number_of_homologs_in_phytopatogens | **54** |
| Checks for homology to mammalian_effectors, and returns the bit score of the best homolog (cut off E.val = 0.01). | homology_to_mamalian.eff_(best_bit_score) | **55** |
| Checks for homology to mammalian_effectors, and returns the number of homologs (cut off E.val = 0.01). | number_of_homologs_in_mamalian.eff | **56** |
| Checks for homology to plants pathogenic bacteria **without** T3SS, using BLASTp, and returns the bit score of the best homolog (cut off E.val = 0.01). | homology_to_plantsPathog**Without**T3ss(best_bit_score) | **57** |
| Checks for homology to plants pathogenic bacteria **without** T3SS, using BLASTp, and returns the number of homologs (cut off E.val = 0.01). | number_of_homologs_in_plantsPathog**Without**T3ss | **58** |
| Checks for homology to plants pathogenic bacteria **with** T3SS, using BLASTp, and returns the bit score of the best homolog (cut off E.val = 0.01). | homology_to_plantsPathog**With**T3ss(best_bit_score) | **59** |
| Checks for homology to plants pathogenic bacteria **with** T3SS, using BLASTp, and returns the number of homologs (cut off E.val = 0.01). | number_of_homologs_in_plantsPathog**With**T3ss | **60** |
| Checks for homology to T3Es of the studied strain (other than the gene itself, if is an effector) , using BLASTp, and returns the bit score of the best homolog (cut off E.val = 0.01). | homology_to_studied_strain_effectors_(best_bit_score) | **61** |
| Checks for homology to T3Es of the studied strain (other than the gene itself, if is an effector) , using BLASTp, and returns the number of homologs (cut off E.val = 0.01). | number_of_homologs_in_studied_strain_effectors | **62** |
| Checks for homology to T3Es of Xanthomonas other than 85-10 strain, using BLASTp, and returns the bit score of the best homolog (cut off E.val = 0.01). | homology_to_Xantho.non8510.eff(best_bit_score) | **63** |
| Checks for homology to T3Es of Xanthomonas other than 85-10 strain, using BLASTp, and returns the number of homologs (cut off E.val = 0.01). | number_of_homologs_in_Xantho.non8510.eff | **64** |
| Computes the genomic distance to the closest T3E. | closest_effector | **65** |
| Computes the number of T3Es in 5 ORFs upstream and downstream to the ORF. | effectors_in_5_ORFs | **66** |
| Computes the number of T3Es in 10 ORFs upstream and downstream to the ORF. | effectors_in_10_ORFs | **67** |
| Computes the number of T3Es in 15 ORFs upstream and downstream to the ORF. | effectors_in_15_ORFs | **68** |
| Computes the number of T3Es in 20 ORFs upstream and downstream to the ORF. | effectors_in_20_ORFs | **69** |
| Computes the number of T3Es in 25 ORFs upstream and downstream to the ORF. | effectors_in_25_ORFs | **70** |
| Computes the number of T3Es in 30 ORFs upstream and downstream to the ORF. | effectors_in_30_ORFs | **71** |
